# Dispersal ability predicts evolutionary success among mammalian carnivores

**DOI:** 10.1101/755207

**Authors:** S. Faurby, L. Werdelin, A. Antonelli

## Abstract

Understanding why some clades contain more species than others is a major challenge in evolutionary biology, and variation in dispersal ability and its connection to diversification rate may be part of the explanation. Several studies have suggested a negative relationship between dispersal capacity and diversification rate among living mammals. However, this pattern may differ when also considering extinct species, given known extinction biases. The colonization of new areas by various lineages may be associated with both diversity increases in those colonising lineages and declines in the lineages already present. Past diversity declines are, however, effectively impossible to infer based on phylogenies of extant taxa, and the underlying process may, therefore, be difficult to determine. Here we produce a novel species-level phylogeny of all known extant and extinct species of the order Carnivora and related extinct groups (1,723 species in total) to show that there is instead a positive relationship between dispersal rate and diversification rate when all extinct species are included. Species that disperse between continents leave more descendant species than non-dispersers, and dispersing species belong to lineages that at the time of dispersal were diversifying faster than the average non-disperser. Our study showcases the importance of combining fossils and phylogenies to better understand evolutionary and biogeographic patterns.

## 1 Introduction

Clades from across the tree of life vary widely in both diversification rate and in the dispersal capacity of the species they comprise, but the extent to which the variation in the two is coupled remains unclear. Among extant vertebrates, a limited number of clades show substantially higher diversification rates than others (1). The fossil record also shows that vertebrate classes vary widely in how many families of equal ages they contain (2). The variation in dispersal capacity among extant vertebrates is equally evident. Some species have colonised nearly the entire world. At one extreme, the range of the wild horse (*Equus ferus*) spanned five continents including northern Africa, the whole of Eurasia and most of the Americas until the end of the last ice age (3). At the other extreme, we find the lemur genus *Eulemur* in Madagascar, where even minor rivers have restricted the migration of individuals sufficiently to result in individual species that are endemic to small areas between neighbouring rivers (4). Interestingly, the clades encompassing these two examples have identical numbers of species (12; *sensu* [3]), at least when the extinct Late Pleistocene species of horses are included (Equidae: *Equus*, *Haringtonhippus,* and *Hippidion*). These two clades also have similar ages (the most recent common ancestor [MRCA] of the lemur genus is ∼4.5 million years [My] old following (4); the MRCA of the horses is ∼6 My following [5]). Taken together, there is thus no universal relationship between dispersal rate and diversification rate.

There are, however, strong arguments for why a relationship between dispersal rate and diversification rate should be expected. A negative correlation between the two rates may be generated under purely neutral models (see e.g. [6]). This is because the *in situ* per area speciation rates would normally increase with decreasing dispersal rates, since populations of poorly dispersing organisms can more readily become isolated from each other and eventually speciate. While a negative relationship is most likely based on neutral models, arguments could also be made for a positive relationship. Increased dispersal rate could potentially increase diversification rate by increasing the total area occupied by the clade, but empirical support for such an effect is limited (7).

While neutral models are thus likely to predict a negative relationship, a positive relationship between diversification and dispersal rates may be the expected outcome of non-neutral models based on interspecific competition (hereafter non-neutral models). A common pattern is a wax- and-wane model with increases followed by decreases in diversity within each clade (e.g. 8, 9). If this is driven by competition, it should produce increased diversity for species with higher dispersal rates, both at the time of diversity increase and the time of diversity decline. During the period of increased diversity, species with a higher dispersal rate would be faster at colonizing all the areas formerly occupied by the species of a clade that they are outcompeting. During their global decline, species with a higher dispersal ability would be able to survive in peripheral regions by escaping direct contact with their competitors.

Although the expectations for non-neutral models have not been explicitly tested, there is anecdotal support based on distribution data, at least when fossils are included. The clearest examples come from the isolated archipelago of New Zealand, where both the Tuatara (*Sphenodon* spp.) and the only known non-flying mammal native to those islands (an unnamed Miocene species often referred to as the Saint Bathans mammal) represent the last remnants of formerly far more diverse and widespread clades (1, 10). There is also evidence of large geographic ranges for some rapidly diversifying lineages, presumably in their expanding phase, such as the Pacific flying foxes (*Pteropus spp*.) (11), but the latter pattern could be generated under both a neutral and a non-neutral model. The expectations from these non-neutral models are also seen in macro-evolutionary analyses of the fossil record. Among mammalian carnivores, there is evidence that the decline in some older clades may be causally related, through increased competition, to the net diversification of younger clades (12, 13).

To reliably tease apart the different mechanisms operating under the neutral and non-neutral models, we may need data on both the diversification rate at the time of dispersal and on the number of descendants each species leaves after dispersal. If diversification rate is increased through increased colonization rate, good dispersers would leave more descendant species than poor dispersers, but they would be unlikely to have a high diversification rate at the time of dispersal in their source area, as discussed above. On the other hand, if non-neutral models prevail, better competitors would both be diversifying faster in their source area and leaving a larger number of descendant species after successful colonization. The required information to test these predictions has, to the best of our knowledge, not yet been assembled.

With few but increasing exceptions (e.g 13, 14), macro-evolutionary studies to date have been based on phylogenies comprised solely of extant species, where the amount of information often makes it impossible to determine if clades are in diversity decline, or show positive yet density-dependent diversification (15). It has even been suggested that estimation of extinction rates relying solely on extant taxa may not be possible (16, 17). Such problems may be avoided for analyses relying solely on fossil data (18). On the other hand, the exclusion of a phylogenetic tree in such analyses usually only allows for comparison of the diversification rates within pre-defined taxonomic entities like families (see e.g. (13)), unlike tree-based analyses where comparisons can be made between any named or unnamed clades. A combined approach based on phylogenetic trees but also incorporating all suitable fossils may, therefore, be optimal for inferring macro-evolutionary patterns (14, 19, 20).

Here we test the relationship between diversification and transcontinental dispersal rates in mammals by combining the advantages of tree-based and fossil-based methods. We build and analyse a complete species-level phylogeny of all extant and extinct species of mammalian carnivores and related extinct groups (Carnivoramorpha, Hyaenodonta, and Oxyaenidae). Our results provide unequivocal evidence that species with high dispersal capacity both had a higher diversification rate at the time of dispersal and left more descendant species than the species that did not disperse between continents. These results suggest that the underlying process is best explained by a non-neutral, competition-driven model.

## 2 Results

### 2.1 Diversity accumulation

We found a nearly continuous increase in diversity across the entire carnivore phylogeny, both globally and within continents, for both phylogenetic and taxonomic diversity (Fig. 1). There were only three main exceptions: 1) Diversity in North America initially peaked at the early Eocene climatic optimum and then decreased in the interval 50 to 45 million years ago (Ma). Given that the majority of species at that time were confined to this continent, a similar decrease was also seen in global diversity. 2) Diversity in both North America and Eurasia decreased over the last ∼10 My. 3) There was a decline in phylogenetic (but not species) diversity in Eurasia between approximately 40 to 35 Ma. Similar results were obtained independently of the length of the time bins being analysed and showed only limited variation across the 100 trees (Fig. 2).

**Figure 1:**
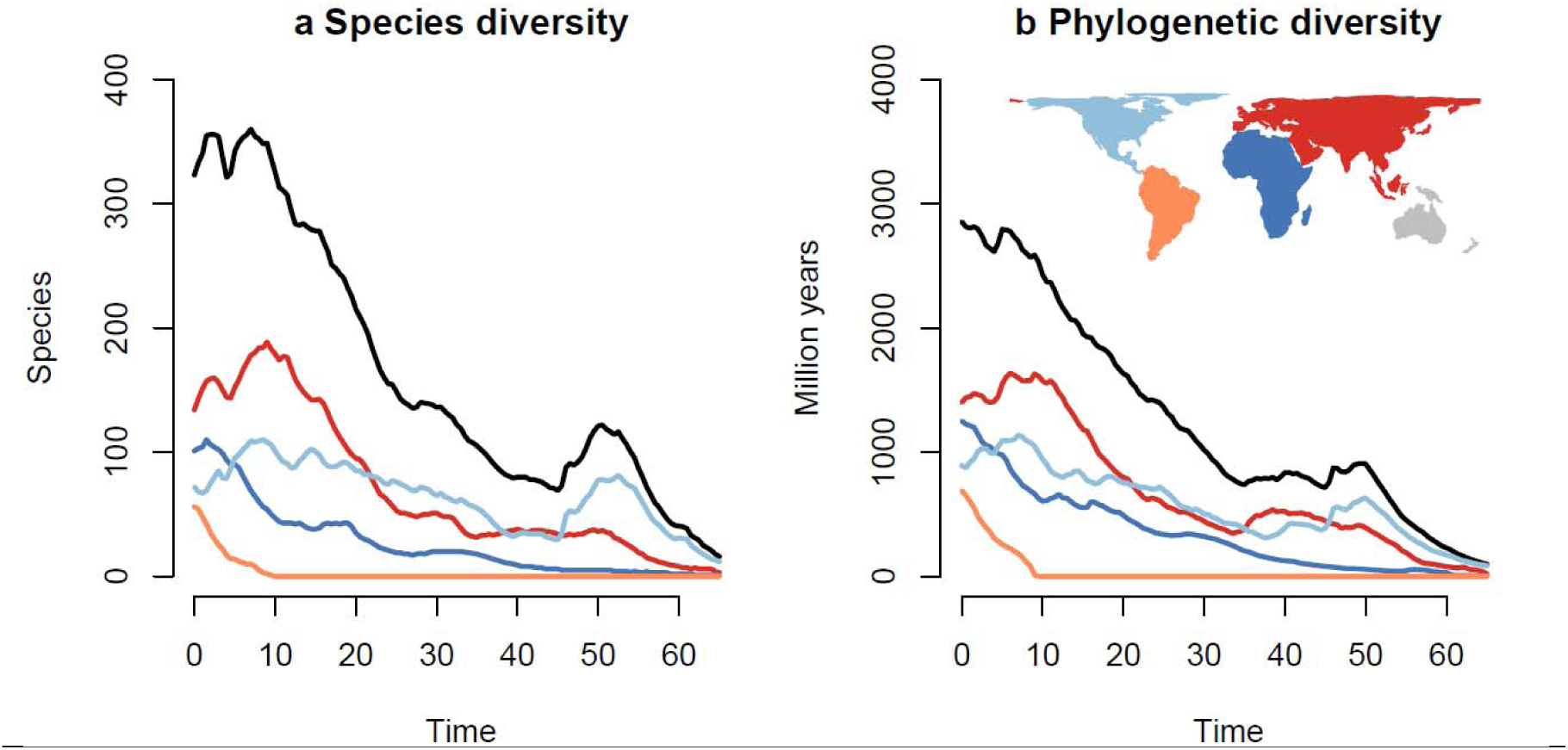
Temporal changes in diversity. Change in species and phylogenetic diversity through time globally (in black) and separately per continent (different colours). Lines represent median values across 100 trees for 0.5-million-year time intervals. The variation between trees and between lengths of the study interval can be seen in Fig. 2.

**Figure 2:**
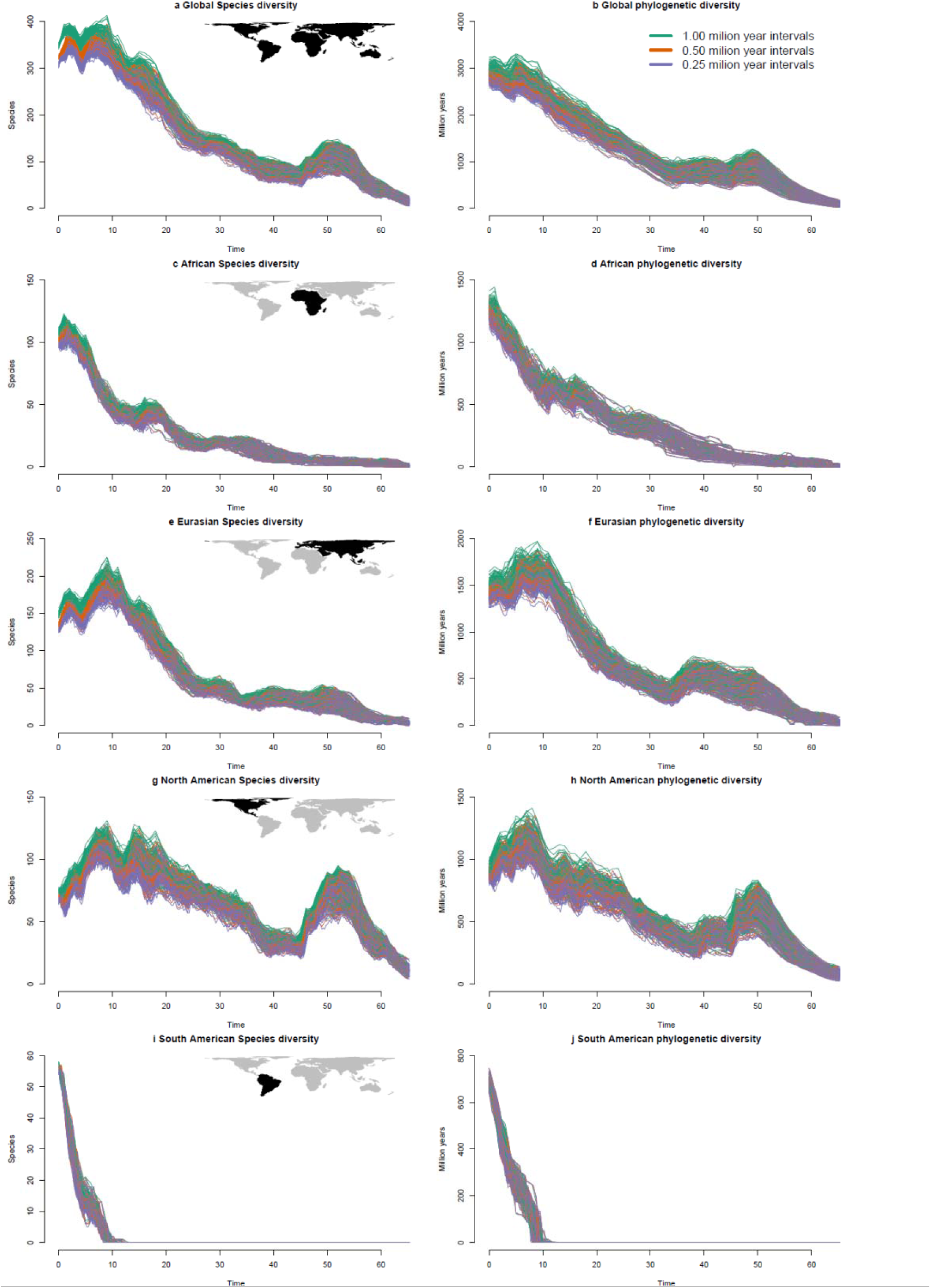
Variation in diversification rate among trees. Plots of diversity through time for 100 trees drawn from the Bayesian phylogenetic analysis. Results for each tree are drawn as separate lines with results for different sample periods shown in different colours. Due to the strong consistency of results between trees and between lengths of the intervals many lines are drawn on top of each other.

These results were based on the assumption of complete sampling. However, the increase in diversity through time could be partially caused by an increase in sampling intensity towards the present. We showed through simulations that this is very unlikely. Our simulations showed very limited effects of incomplete sampling on the observed patterns when using empirically derived sampling intensities (Figure S1).

### 2.1 Higher evolutionary success of dispersers

We estimated the evolutionary success of dispersers using two novel metrics, which we refer to as *pre-dispersal success* and *post-dispersal success* (Fig. 3). *Pre-dispersal success* measures the diversification rate of lineages at the time of their dispersal, whereas *post-dispersal success* measures how many species the dispersing lineages diversify into.

**Figure 3:**
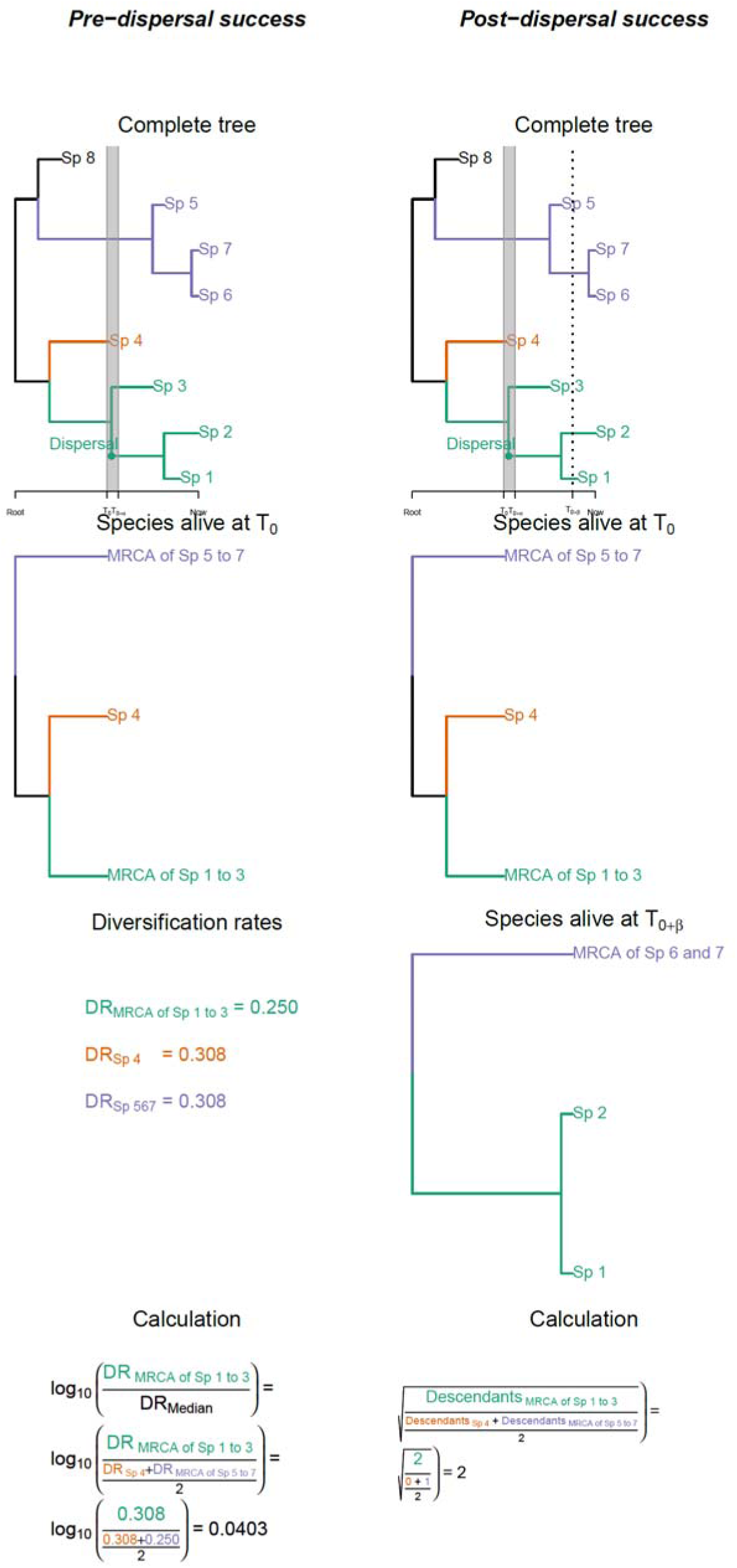
Estimation of dispersal success. In the example above, one dispersal event happened within the interval from T_0_ and T_0+α_ represents the time interval, which in our case was 0.25, 0.5 or 1.0 million years (My).. where α Both *Pre-dispersal success* and *Post-dispersal success* are calculated on the same tree. The panels on the right illustrate how *Pre-dispersal success* is calculated. In order to improve understanding, we have used the same colours on each panel to show corresponding parts of trees or equations. The two first panels show the entire tree, and a tree of just the species alive at time T_0_. The third panel shows DR rates estimated as in (44) for the species alive at T_0_. The fourth shows the calculation of *Pre-dispersal success*, which we define as the logarithm to the ratio between the DR of the dispersing lineage and the median of the remaining lineages (see the methods section for details). The panels on the left illustrate how *Post-dispersal success* is calculated. In order to improve understanding, we have used the same colours on each panel to show corresponding parts of trees or equations. The first panel shows the entire tree. This is identical to the tree for *Pre-dispersal success* except that a stippled line has been added at time T_0+β_. β here represents a pre-defined length of time (in our case 3, 5 or 7 My). *Post-dispersal success* compared the trees at T_0_ and T_0+β_ and how many descendant species, each species alive at T_0_ has diversified into. The next two panels show the trees of the species alive at T_0_ and T_0+_ respectively. The last panel illustrates the calculation of *Post-*β *dispersal success*, which we defined as the square root of the ratio between the number of descendants of the dispersing lineage and the mean number of descendants for any lineage from time T_0_ alive at time T_0+_. Note that we here only look at the descendants alive at this time point and not all descendants. This, for instance, means that the taxon “*MRCA of Sp 5 to 7*” has only diversified into one species at time T_0+β_ (“*MRCA of Sp 6 and 7*”) because Sp 5 is already extinct by then and the split between Sp 6 and Sp 7 happens at a later stage (see the methods section for details).

#### 2.2.1 Pre-dispersal success

Our analyses of *pre-dispersal success* suggest that the dispersing species belong to clades that, at the time of dispersal, were diversifying faster than non-dispersers (Tables 1, S1-S2). This pattern was observed irrespective of whether comparisons were to all species alive in the time interval, or only to species occurring on the source continents in the time interval (which we refer to as *global* and *continental pre-dispersal success*). We estimated dispersal within time bins rather than in continuous time, but the results were independent of the length of these bins. The best model for *continental pre-dispersal success* showed a difference in success between dispersers and non-dispersers depending on the target continent. In particular, the model showed a substantially smaller difference between dispersers and non-dispersers for species colonizing South America. The best model for *global pre-dispersal success* showed temporal variation, where the difference in success between dispersing and remaining lineages was smaller for older dispersal events. In both cases, however, both models had lower AIC than the model without any spatial or temporal variation. Thus, the two analyses only disagreed on whether spatial or temporal variation was most important.

**Table 1:** Pre-dispersal success. Pre-dispersal success was estimated as the logarithm of the ratio between the diversification rate of the dispersing lineage dispersers and the median of the dispersal rates of the remaining species alive at the time of dispersal (see Fig. 3). Separate analyses were conducted comparing the dispersers with either all species alive (*global pre-dispersal success*) or only the species alive in the source continent (*continental pre-dispersal success*). The p-values for node age and for global rate are the probability of the estimate in question being greater than 0. For models with different patterns depending on continental source and target, the p-value is based on the probability of being different from the estimated global rate. This table only lists the results for the simplest model, and models preferred by AIC for either global or *continental pre-dispersal success*. The results from the remaining models are provided in the supplementary material (Tables S1–S2). Values are only given for time intervals of 0.5 million years, but results are similar for the other two intervals (Tables S1–S2).

|  | *Continental pre-dispersal success* | *Global pre-dispersal success* |
| --- | --- | --- |
| Simplest model |  |  |
| Global | 0.0835 (0.0194)*** | 0.1205 (0.0205)*** |
| ΔAIC | 5.973 | 6.178 |
| Temporal variation |  |  |
| Global | 0.1122 (0.0259)*** | 0.1636 (0.0272)*** |
| Age before present | -0.0018 (0.0012)‘ | -0.0027 (0.0013)* |
| ΔAIC | 4.079 | 0.000 |
| Variation between target continent |  |  |
| To Africa | 0.1160 (0.0373) | 0.1468 (0.0372) |
| To Eurasia | 0.0453 (0.0343) | 0.1050 (0.0362) |
| To North America | 0.1260 (0.0360) | 0.1420 (0.0370) |
| To South America | 0.0017 (0.0503) | 0.0423 (0.0528) |
| ΔAIC | 0.000 | 5.873 |
| ‘ 0.10>P>0.05 * 0.05>P>0.01 ** 0.01>P>0.001 *** 0.001>P |  |  |

**Table 2:** Post.dispersal success. Post-dispersal success is estimated as the square root of the ratio between the number of descendants for each dispersing lineage alive 5 million years (My) after dispersal divided by the mean number of descendants for any species alive at the time of dispersal (see Fig. 3). Separate analyses were conducted comparing the dispersers with either all species alive then (*global post-dispersal success*) or only the species alive in the target continent (*continental post-dispersal success*). P-values for node age and for global rate are the probability of the estimate in question being higher than 1 (the null expectation). For models with different patterns depending on continental source and target, the p-value is based on the probability of being different from the estimated global rate. Values are only given for time intervals of 0.5 My and only based on the number of species alive after 5 My, but results are similar for time intervals of 0.25 and 1.00 and after 3 or 7 My (Tables S5–S6). This table only lists the results for the simplest model, and models preferred by AIC for either *global* or *continental post-dispersal success*. The results from the remaining models are provided as supplementary material (Tables S5–S6).

|  | *Continental post-dispersal success* | *Global post-dispersal success* |
| --- | --- | --- |
| Simplest model |  |  |
| Global | 1.4433 (0.0730)<sup>***</sup> | 1.4606 (0.0730)<sup>***</sup> |
| ΔAIC | 5.792 | 0.000 |
| Variation between source and target continent |  |  |
| Africa to Eurasia | 1.2666 (0.1727) | 1.3557 (0.1777) |
| Eurasia to Africa | 1.3607 (0.1206) | 1.3991 (0.1184) |
| Eurasia to North America | 1.6677 (0.1282)<sup>‘</sup> | 1.5767 (0.1243) |
| North America to Eurasia | 1.4238 (0.1487) | 1.4572 (0.1428) |
| North to South America | 1.1263 (0.2925) | 1.4005 (0.2835) |
| ΔAIC | 0.000 | 1.685 |
| <sup>‘</sup> 0.10>P>0.05 * 0.05>P>0.01 ** 0.01>P>0.001 *** 0.001>P |  |  |

Our analyses assumed complete sampling of all extinct species but we tested the consequences of this assumption through simulations. For this, we modelled scenarios of no difference in diversification patterns between dispersers and non-dispersers and assessed if spatial patterns in sampling would create a false signal with such a difference. These simulations showed that the patterns of *pre-dispersal success* were not caused by incomplete sampling (Table S3-S4). The simulations of *global pre-dispersal success* found no significant difference in success between dispersers and non-dispersers and found no support for any spatial or temporal variation in the difference between dispersers and non-dispersers. The simulations of *continental pre-dispersal success* also found no significant difference in success between dispersers and non-dispersers. They did, however, recover weak support based on AIC for models with temporal, but not spatial, variation in the difference between dispersers and non-dispersers. Even then, the estimated effect size for temporal variation was not significantly different from zero. If sampling effort did have an effect on spatial or temporal variation in our results, the effect size must have been minimal.

#### 3.2.2 Post-dispersal success

Our results clearly demonstrate that dispersing lineages leave more descendant lineages than lineages that remain within the source continents (Table 2). This applies to both *continental* and *global post-dispersal success* (i.e. comparisons to all other species on the target continent or all species alive at the time of dispersal). This pattern – that dispersers leave more descendant lineages – remained constant irrespective of the length of the analysed time bin and how long after the dispersal the number of descendant species was counted (Tables S5-S6). The model with spatial variation was, however, only supported for *continental* (but not *global*) *post-dispersal success* and similar, although weaker, support for spatial variation was also recovered when we simulated incomplete sampling (Table S7). We, therefore, focus our discussion solely on the strong evidence for higher *post-dispersal success* rather than on any more detailed spatial or temporal patterns regarding the magnitude of this success.

## 3 Discussion

Our results unequivocally show a positive correlation between diversification rate and dispersal in carnivores. The analyses are based on the first species-level phylogeny of carnivores that includes all suitable fossils and all extant species. These results contradict an expected neutral pattern of a trade-off between diversification rate and dispersal, as has been suggested based on analyses of contemporary mammals (6).

### 3.1 Diversity accumulation

The occasional periods of diversity decline detected by our analyses lend biological credence to our results, since a monotonous increase in diversity could point towards a pattern driven by insufficient fossil information. All three declines detected clearly match previous knowledge. The first two declines were likely climatically driven. If carnivores throughout their history have had lower diversity in higher latitudes, similar to what we see today (21), we should expect to see diversity declines during times of global temperature decline, particularly in North America and Eurasia, which have the highest proportions of non-tropical areas. In this regard, the first early decline in North America coincides with a period of Eocene cooling (22) and is temporally similar to a time period recently found to have a low overall mammalian diversity in North America (23). Secondly, the declines in North America and Eurasia during the last 10 My may be explained by the overall climatic cooling during this time period (24).

In contrast, the third decline in phylogenetic diversity in Eurasia may not have been climatically driven. Instead, it probably reflects the so-called ‘Grande Coupure’, where the formerly isolated European fauna was replaced by an immigrant Asian fauna (e.g. 25). The cause of the Grande Coupure is not entirely known (25) but if it represents biotic replacement driven by competition, which is one of the hypotheses (25), it may indicate that such processes are important for carnivores in general.

### 3.2 Neutral or non-neutral models

Our results for *pre-dispersal success* and *post-dispersal success* clearly suggest that dispersal rate and diversification rate are linked in carnivores. As previously noted, the pattern for *post-dispersal success* could be the result of either neutral or non-neutral models or both, but the higher *pre-dispersal success* among good dispersers is only expected under the non-neutral models.

Support for a non-neutral model is further evident in the temporal variation in *pre-dispersal success.* We find an increasing effect for recent dispersals, where the number of free niches would be expected to be lower. This matches the expectations of this model – the non-neutral model would only generate a relationship between *pre-dispersal success* for dispersers and non-dispersers if dispersal to other continents required the displacement of lineages already there. There should be smaller differences if there are free niches open to any coloniser. This non-neutral model, therefore, contrasts with the frequently found priority effects in community ecology where the first coloniser is nearly always more successful (26).

The spatial patterns also support non-neutral models (Table 1). We find elevated *pre-dispersal success* for dispersers to North America and a near-random pattern for dispersers to South America, which matches our expectations. Under non-neutral models, we only expect elevated *pre-dispersal success* for dispersers if these are invading already occupied niches. The South American continent lacked placental carnivores until the mid-Miocene (27), and all invading carnivores would have initially encountered empty ecological niches. The pattern of elevated *pre-dispersal success* in North America is expected because our analyses suggest that carnivores originated in North America. Our results also show that the clade has consistently been highly diverse in that continent (Fig. 1). This North American origin is clear from Oxyaenidae, which is one of the three earliest diverging clades we analyze (28). The origin of the two other clades (Carnivoramorpha and Hyaenodonta) has previously been considered to be Eurasian or African, but even studies suggesting a non-North American origin for these clades have suggested extremely rapid dispersal to, and substantial diversification within, North America (29, 30).

Our results partially contradict earlier work which suggested that dispersal from North America to Eurasia – but not in the opposite direction – was associated with high diversification rate (31). Our results suggest the opposite and we found both higher increases in both *pre-dispersal* and *post-dispersal success* for dispersers to North America compared to dispersers to the other continents (Tables 1–2). The reason for this difference may partly be a function of the non-phylogenetic approach of Pires et al (31), meaning that the different outcomes of multiple dispersals within the same family could not be distinguished.

As a minor point of uncertainty, we note that we treat the carnivore niche in South America as unoccupied, although it was occupied prior to the arrival of the placental carnivores by the Sparassodonta (Metatheria; sister group to marsupials). It is still unknown if the Sparassodonta went extinct independently of the arrival of placental carnivores, or if they were driven to extinction by competitive replacement (32). Non-placental lineages seem, however, to be inherently inferior competitors to placental carnivores, likely due to effects related to their lack of deciduous teeth (33, 34, 35). It, therefore, seems plausible that if the Sparassodonta were still extant when the carnivores arrived then any member of the group that arrived there may have been able to outcompete them.

### 3.3 Implications

Our results suggest a model of consistent competitive replacement among carnivore clades, although the generality of the observed pattern remains unclear. The methodology we employed was possible because carnivores have a well-understood fossil history, which is why the effects of incomplete sampling were deemed minor (Tables S3–S4, S7; Fig. S1). Furthermore, carnivores are a particularly useful group to study for this purpose because there is strong phylogenetic conservatism in their niche, with few other taxa competing with them. They are thus nearly a monophyletic ecological guild, although there are exceptions to this; some species within the group, such as the giant and red pandas, for instance, are predominantly herbivorous (3). Until the Eocene, carnivores shared the carnivorous niche with other mammalian species of uncertain placement, such as Arctocyonidae or Cimolestidae (36), but even then, carnivores plausibly comprised the majority of the guild (12). Arguably, the only other large monophyletic mammalian group that behaves like an ecological guild is bats, but they are noteworthy for having a particularly scarce fossil record among mammals (37). It may therefore not be possible at present to directly replicate our analyses using other clades and thereby directly test the generality of the patterns reported here.

Despite the difficulties in applying our methodology to other clades, we find it unlikely that the patterns we report here would be taxon-specific. Taxon specificity could have explained why our results appear to run counter to some studies showing lower diversification rates for species with higher dispersal rates (6, 7). We think, however, that the apparent differences are instead a consequence of the taxonomic scale of the analyses and the inclusion or exclusion of fossil taxa. A lower diversification rate in highly dispersing lineages has been recovered in analyses at the subfamily or family level, whereas our estimates here are conducted at the species level. Direct competition may be strongest between the most closely related species in the case of mammals (38), which is also what would be expected for any traits with phylogenetic conservatism. This would explain the apparent conflict between this and earlier analyses of mammals. Earlier analyses at the subfamily level have suggested a negative relationship between dispersal and diversification rate in mammals (6), whereas in a species-level analysis we here recover a positive relationship between the two.

We are not implying that competition is not frequent between distantly related taxa, as is increasingly being acknowledged (39). However, competition may be expected to be linked to physiological or morphological traits, which are generally more similar for closely related species. There are many examples of convergent evolution within mammals (e.g. 40, 41), but even so, recently diverged species will be more similar than a random pair of species under most evolutionary models. They must, therefore, be expected to rely more on the same resources and the same environmental conditions than random members of a larger clade.

If the pattern we recover is driven by non-neutral biotic interactions, it may only be observable because we included fossil species in our analyses. When species are driven to extinction by other species it may be on too fast a time scale for us to see it clearly based only on extant species in their native ranges. The only clear contemporary evidence for biologically caused extinctions or declines comes from the invasion biology literature (e.g. 42). Competition-driven extinctions could leave signals on the phylogenies of the extant species but, as we noted in the introduction, such declines may be extremely difficult to detect based on phylogenies of extant species only (15, 16, 17). Even in the few situations where declines can be detected, methods solely relying on contemporary species can logically only give a signal if the declining lineages still have at least one extant species. Some of the clearest cases of clade competition, such as the bone-crushing dogs (Borophaginae) being driven to extinction by related and extant modern dogs (Caninae) (13), are thus impossible to infer without fossils.

In summary, our analysis of a novel species-level phylogeny of all extant and extinct carnivores shows that: 1) lineages that disperse between continents are generally those that diversify more quickly; and 2) lineages that colonise new continents leave more descendant species than lineages already present there. These results are only likely to have emerged because we combined fossil and phylogenetic information, highlighting the need to incorporate both sources of information whenever possible.

## 4 Methods

### 4.1 Method summary

We analysed all extant and extinct species of mammalian carnivores and related extinct groups (Carnivoramorpha, Hyaenodonta, and Oxyaenidae). Herein we refer to this entire clade as ‘carnivores’. We revisited the taxonomy of all fossil and extant members of the group and accepted, 1723 species (314 of which are extant). We based our analyses on records in the Paleobiology Database (PBDB; https://paleobiodb.org/) and the New and Old Worlds Database of fossil mammals (http://www.helsinki.fi/science/now/; NOW), but supplemented these with data from the original literature for 128 species that we consider valid but which, at least when we were collecting data, lacked any records in either of the two databases.

We constructed the phylogeny of all extant and extinct species of carnivores using a tip dating approach under a fossilised birth-death model in MrBayes 3.2 (43). We did this in a two-step procedure combining a backbone tree with a number of smaller phylogenies at lower taxonomic levels. This procedure is similar to that used to construct phylogenies focusing on other large clades (e.g. 44, 45), but it has previously only been used to generate all-taxon phylogenies of all extant species within a clade. This is the first time it has been expanded to include all extant and extinct species within the focal clade. The placement of species without genetic or morphological data was facilitated by a number of constraints based on taxonomy and suggested relationships from taxonomic treatments. These trees only give species origination time, but the information from these was combined with extinction times generated by the Bayesian program PyRate (46), which estimated likely extinction times based on the temporal distribution of all known records of each species. The resulting phylogeny is attached as appendix 1 giving 1000 trees from the posterior distribution of trees We included pinnipeds to improve the usability of the phylogeny for other researchers, but here we discarded them for all analyses for this paper due to our focus on terrestrial species.

We inferred the ancestral areas of all nodes based on a DEC (dispersal-extinction-cladogenesis) model in BiogeoBEARS (47). We used a DEC rather than the DEC +j model since the underlying mathematical properties of the DEC +j model have been questioned (48). Following the estimation of ancestral areas for all nodes, we inferred dispersal events and times along branches.

We assessed changes in global and continental diversity by plotting species and phylogenetic diversity (49) (i.e. the sum of branch lengths). Following this, we analysed the evolutionary success (estimated as their diversification rate) of dispersers at the time of their dispersal (*pre-dispersal success*) and the number of descendant species they left behind after a set time (*post-dispersal success*).

The analyses related to phylogenetic diversity and diversification rate are only meaningfully interpretable for ultrametric trees. For simplicity, extinction was therefore dealt with in time intervals rather than in continuous time and on trees sliced at various ages, only counting the species (internal or external branches) extant at that point in time. Hence, when using 0.5 My time intervals, two species that went extinct 1.2 and 1.4 Ma were assumed to have survived until 1.0 Ma and would both be included as extant for a tree sliced at 1.0 Ma. To test the effect of this procedure on the results, all analyses were conducted with time intervals of 0.25, 0.5 or 1.0 My duration.

We estimated *pre-dispersal success* based on diversification rate (DR) (45). We sliced the tree at the end of each time interval throughout the Cenozoic and calculated the DR of all lineages alive at that time. For all intercontinental dispersal events occurring in the following time interval, we then identified the lineage at the beginning of the interval that would evolve into the disperser during the interval. This could be either the same species or one of its ancestors, which would be the case for founder speciation occurring within the time interval. We then calculated the logarithm of the ratio between the diversification rate of the disperser and the median diversification rate for either all lineages alive in the time interval (*global pre-dispersal success*) or the subset of these that was found on the same continent the disperser originated in (*continental pre-dispersal success*). This calculation is outlined in Fig. 3.

We estimated *post-dispersal success* by comparing the tree at the time before dispersal with the tree sliced a number of million years afterwards. For each lineage alive at the first time interval, we identified how many species it had diversified into a few million years later (which was 0 if the lineage had gone extinct in the meantime). We then calculated the ratio between the number of species in the dispersing lineage and the mean for either all other species (*global post-dispersal success*) or all species from the continent dispersed to (*continental post-dispersal success*). In both cases, we square-root-transformed the *post-dispersal success* to improve normality. Separate analyses of *post-dispersal success* were conducted on trees sliced after 3, 5, and 7 My.

Although we have included all known species, an unknown number of extinct species may be missing from the fossil record, which may influence our results, especially since the fraction of missing species is likely to vary in time and space. In order to understand the influence of missing taxa on our results, we, therefore, simulated a number of random phylogenies. We then simulated incomplete sampling on those phylogenies based on spatial and temporal sampling effort estimated in the PyRate analyses described above. Following this, we repeated all analyses described above on both the full and the sampled phylogenies in order to directly measure the effect of incomplete sampling on our results.

A detailed explanation of all steps can be found in the supplementary materials and methods.

## Supporting information

A description of our treatment of all the individual fossil records

The produced phylogeny of all carnivores

## Supplementary Materials

Supplementary materials and methods

Tables S1-S9.

Figures S1-S2.

Appendix 1: The produced phylogeny of all carnivores

Appendix 2. A description of our treatment of all the individual fossil records

## Acknowledgments

### Funding

Funding for this work was provided through a Wallenberg Academy Fellowship from the Knut and Alice Wallenberg Foundation, by the Swedish Research Council (B0569601, 2015-04587, and 2017-03862), the Swedish Foundation for Strategic Research and the Royal Botanic Gardens, Kew. Author contributions: SF, LW, and AA designed the study. SF and LW performed the taxonomic cleaning for all records. SF conducted all analyses. SF led the writing of the paper with input from LW and AA. All authors approved the submitted version.

### Competing interests

Authors declare no competing interests.

### Data and materials availability

The produced phylogeny is added as an appendix. All scripts used for running individual analyses are available from the authors upon request.

## Supplementary materials

### 5 Supplementary materials and methods

#### 5.1 Phylogeny

##### 5.1.1 Input data

We downloaded all records of carnivores identified at least to genus level from the Paleobiology Database (PBDB; https://paleobiodb.org/) on October 10, 2016 and the New and Old Worlds Database of fossil mammals New and Old Worlds Database of fossil mammals (http://www.helsinki.fi/science/now/; NOW), on September 30, 2016. We defined the focal clade as Carnivoramorpha (Carnivora and Miacoidea) + Creodonta (Hyaenodonta and Oxyaenidae). Hereafter, we refer to this entire clade as ‘carnivores’ and use the term ‘Carnivora’ when referring exclusively to the extant order.

We revisited the taxonomy of all named species to generate a consistent list matching current knowledge of extinct mammals, as well as the taxonomy of species surviving at least until the Late Pleistocene (hereafter ‘extant carnivores’). For the latter, we followed the taxonomy of the Phylacine V 1.2 database (3). Phylacine follows the International Union for Conservation of Nature (IUCN) version 2016-3 for extant species and extinct species with extinction dates post 1500 AD, and an updated version of the database of Faurby and Svenning (44) for species that went extinct between the Late Pleistocene and 1500 AD. The resulting dataset had 7,551 useable records from NOW (6,285 records assigned to a species and 1,266 only to genus, while 94 records from the database were not assignable to any of the genera we accept and were therefore excluded) and 7,984 useable records from PBDB (6,755 records assigned to species and 1,229 only to genus, while 123 were not assignable to any of our genera and therefore excluded).

Our combined dataset consisted of 1,723 species after cleaning, 314 of which are extant. Among the species that went extinct prior to the Late Pleistocene, 631 were included in both NOW and PBDB, 382 only had records in the NOW Database, and 268 only had records in PBDB. An additional 128 species were manually added since they were not included in either of the databases at the time of original download (October 10, 2016, and September 30, 2016 for the PBDB and NOW databases, respectively; see Table S8). We included a few forms as separate species entities although not formally described as such. Five genera had records only identified at the genus level from North America. For our analyses, we treat these as distinct species, such as “*Parailurus* NorthAmerica”.

Although species designation for these has not formally been made, morphological differences have generally been noted (50), which makes species designation plausible. Finally, we treat the records of two small species (*Palaeogale minuta* and *Palaeogale sectoria*) from North America and Eurasia as distinct continental endemics, since both persist on both continents for many million years and it seems biologically implausible for them to maintain population coherence (i.e., gene flow) during that time interval. A full breakdown of records by database can be found in the attached Excel spreadsheet ‘Database summary’.

##### 5.1.2 Phylogenetic and dating analyses

The phylogeny of all extant and extinct species of carnivores was constructed using a tip dating approach under a fossilised birth-death model in MrBayes 3.2 (43). The phylogeny was created by combining a backbone phylogeny with 17 smaller phylogenies (for Amphicyonidae, Barbourofelidae, Canidae, Eupleridae, Felidae, Herpestidae, Hyaenodontidae, Hyaenidae, Mustelidae, Nimravidae, Oxyaenidae, *Palaeogale*, Percrocutidae, Pinnipedia, Ursidae, Viverravidae and Viverridae). This procedure requires that all the smaller phylogenies have a known number of species. For two families (Miacidae and Stenoplesictidae) that were not constrained to be monophyletic (see next section), we, therefore, included all species in the backbone phylogeny. Two chains were run for both the backbone and the 17 smaller phylogenies until the average standard deviation of split frequencies was lower than 0.03, but for a minimum of 10 million generations. The analyses were further inspected with Tracer 1.6 (51) to ensure that the Effective Sample Size of the post burn-in for the overall model tree likelihood for the two chains combined was at least 200.

The priors for the analyses were based on earlier tip dating analyses across all mammals from Ronquist et al (52). In particular, we set a uniform prior of the root between 56.3 and 88.0 Ma, representing the span between the oldest fossil in the database and the estimated divergence time between the MRCA of Hyaenodontidae and Carnivora and their combined outgroup, following Ronquist et al. (52). We further penalised long ‘ghost lineages’ (i.e. lineages existing for very extended time periods without leaving any fossil evidence) in all analyses using the prior “prset fossilizationpr = beta(100, 1)” as also suggested by Ronquist et al (52). For all phylogenetic reconstructions, we assumed that all species are included in the analyses. While we have included all known species in the phylogeny, we acknowledge that a fraction of undescribed species must be missing, but this fraction cannot be reliably estimated with any available method.

The backbone phylogeny was constructed based on morphological data for extinct clades and a combination of morphological and genetic data for extant ones. All morphological data were analysed under an MK-model conditioned to only include variable sites and incorporating gamma rate heterogeneity. The morphological data were based on the matrix by Wesley-Hunt and Flynn (53) but supplemented by numerous studies (see Table S9). We added new data for four key taxa representing taxonomic groupings not included in previous analyses (*Percrocuta* sp, *Ginsburgsmilus napakensis*, *Barbourofelis* sp, and *Oxyaena forcipata*) (all four coded by LW). This morphological matrix was supplemented with genetic data from Meredith et al (54). To facilitate the merging of the smaller phylogenies with the backbone phylogeny, we added the oldest known species of each family (in this and all other cases we use the ages as listed in the original data source) to the backbone analysis, with all characters coded as missing data. The 17 smaller phylogenies were for the most part constructed based on one or more morphological data matrices, with genetic data also included for the extant families. Depending on the family, the genetic data were either based almost entirely on a single source although with supplemented searches for missing data of missing species from NCBI, or on NCBI searches for each species (a list of sources can again be found in Table S9).

We made a number of modifications to the nexus files before running MrBayes. We set the best nucleotide substitution model and partition scheme for the genetic data as the optimal one based on AIC as inferred by Partitionfinder 1.1 (55) for both the overall and the smaller phylogenies. We set the age of all fossil taxa and all extant species without genetic data to the age of the oldest known record. Six of the species not included in either NOW or PBDB (*Amphicynodon brachyrostris*, *Amphicynodon cephalogalinus*, *Amphicynodon chardini*, *Amphicynodon crassirostris*, *Phoberogale mino*r and *Filholictis filholi*) come from undated deposits. The first five were given uniform priors between the age of the oldest and youngest species in the genus, whereas the sixth (*Filholictis filholi*) is from a monospecific genus and was given a uniform prior between the youngest and oldest member of the subfamily. For all genera constrained to be monophyletic (see next section), we set the minimum age of the genus to the age of the oldest record in the genus (whether identified to species or not). We did this using a uniform prior on the node age, with a maximum age equal to the rootage of the family. We set the prior on the age of all tips representing fossil species and all extant species lacking genetic data as a uniform prior ranging between the minimum and maximum ages of the oldest known fossil of the species. These are intended to inform the origination time for the species lacking genetic data. For all later analyses, they are treated in the same way as the extant taxa.

We merged the smaller trees with the backbone phylogeny while keeping the dating information for both sets, as described below. For the 17 smaller phylogenies, we set a uniform prior of the rootage between the oldest known fossil in the group and the upper 95% HPD (Highest Posterior Density) for the stem age of the group from the backbone phylogeny. The backbone and the smaller phylogenies were merged so that there was the same correlation between the stem and crown age as between the age of the stem age and the next internal branch in the phylogeny. That is, since Felidae and Barbourofelidae are sister families, the phylogenies were merged so the correlation between the stem and rootage of Felidae is the same as the correlation between the stem age of Felidae and the age of the MRCA (Most Recent Common Ancestor) of Felidae and Barbourofelidae.

For some trees, the resulting family-level clades had crown ages slightly older than the stem ages of the overall tree, and we, therefore, needed to recalibrate the family level trees to avoid negative branch lengths. This was done for all problematic trees (i.e. family level trees with crown ages slightly older than the stem ages from the backbone tree) so that all branch lengths in the new tree were proportional to the branches in the original tree, and the rootage was equal to the stem age of the backbone tree minus 0.01.

##### 5.1.3 Constraints

Similar to the assumption of other complete phylogenies, where a number of species lack genetic or morphological data (e.g. 44, 45), we assumed taxonomic clades to be monophyletic unless there is good evidence against it. This meant that we carefully inspected the paleontological literature regarding each taxonomic unit to judge if they are generally understood to comprise monophyletic entities (all taxonomic constraints are shown in the attached Excel spreadsheet ‘Database summary’).

At the highest level Carnivoramorpha, Hyaenodonta, and Oxyaenidae were each constrained to be monophyletic. Within Carnivoramorpha, Carnivora and Viverravidae, but not Miacidae (which is a paraphyletic assembly; see e.g. 53), were constrained to be monophyletic. Within Carnivora, all taxa were constrained to be either Caniformia or Feliformia. Within Caniformia, we constrained the monophyly of Amphicyonidae, Canidae, Musteloidea, Pinnipedia, and Ursidae and assumed *Lycophocyon hutchisoni* to be outside any of the major lineages. Within Feliformia we constrained the monophyly of Barbourofelidae, Eupleridae, Felidae, Herpestidae, Hyaenidae, Nimravidae, Percrocutidae, Prionodontidae, and Viverridae, but not Stenoplesictidae (which again is generally considered a paraphyletic assemblage; see e.g. 56). This means that we assumed that each genus normally assigned to Stenoplesictidae, as well as *Palaeogale*, was outside any of the major lineages listed above. We further constrained Percrocutidae as sister to Hyaenidae following (57) and Felidae as sister to Barbourofelidae following (58). The morphological dataset started by Wesley-Hunt and Flynn (53) was designed to determine the relatedness between basal taxa and on its own (i.e. without genetic data added) produces improbable relationships between more derived members of extant families within Carnivora. (53). Both Percrocutidae and Barbourofelidae are only known from Miocene fossils, meaning that the morphological matrix may be suboptimal to infer their placement, but unlike the extant families, their relationship cannot be inferred by adding genetic data to the analysis.

At a lower level, we constrained most subfamilies and genera to be monophyletic, but deviated from this in a number of cases for three main reasons: 1) Some of the earliest described genera within families and subfamilies, e.g. *Lutra* for otters, have served as waste-baskets for a number of frequently poorly-defined fossil taxa (59). Many fossil forms within such genera, as well as other poorly known taxa, were therefore allowed to be placed freely within the family or subfamily instead of being constrained to their genera; 2) Other species of uncertain phylogenetic placement belong to distinct genera, but are rarely included in the newer taxonomic treatments and were therefore not constrained to be within otherwise constrained subfamilies or tribes; and 3) Taxonomy does not always imply genus-level monophyly. For some taxa, there is evidence that named genera are nested within other named genera and we, therefore, allowed such nesting when supported. For example, this is the case for *Neovison* (the American mink and the extinct sea mink), which phylogenetically may be nested within *Mustela* (weasels) (60).

In addition to these taxonomic constraints, we also employed a number of constraints based on stated likely relationships in taxonomical treatments. Finally, we employed a number of biogeographical constraints within lineages or species and often enforced that there would only be a single intercontinental dispersal within a lineage unless there are data to suggest otherwise. A full breakdown of family, subfamily and genus level constraints can be found as part of appendix 2, which contains information on all fossil records and our treatment of them, while a full list of additional constraints and the relevant sources can be found in Table S10.

##### 5.1.4. Extinction times

The procedure described above only gives the origination times of all lineages, but not the extinction times. To estimate the actual extinction times for all taxa we used the Bayesian program PyRate (45). We did this independently for each continent, which means that we treat a species occurring on multiple continents as two distinct populations that may go extinct independently of each other, rather than as a coherent group of sub-populations with ongoing gene flow.

Firstly, we combined fossil records from NOW and PBDB, keeping as many records as possible while avoiding duplicate records. For each species, we initially accepted all records in either NOW or PBDB (whichever had most records of the species in question). We then examined all records of the other database for the same species, one by one, to assess if they were clearly distinct (in which case they were added) or potential duplicates (in which case they were not). If the latitude and or longitude rounded to the nearest degree was different from all records already accepted, and/or if the age of the record was non-overlapping with accepted records with the same latitude and longitude, we added the record. If there was only one record already accepted with identical latitude and longitude and overlapping age, we considered the old and new ones to be potential duplicates and retained the one with the most precise dating. However, we discarded the new record if there was more than one already accepted record with the identical latitude and longitude and overlapping age.

Secondly, we ran PyRate analyses separately for records from South America, North America, Asia, Europe, and Africa, with an additional separate analysis for Pinnipeds (since marine species may have different fossilization potential than terrestrial ones). Sampling intensity (i.e. the product of the number of specimens fossilizing and the fraction of fossils that are identified and placed in the available databases) is a vital component determining how long after the youngest record the true extinction time is likely to have been. Sampling intensity may vary between continents and between marine and terrestrial species. For each continent (and for pinnipeds) we ran 20 separate PyRate analyses, further allowing for variable sampling intensity in each epoch, for example as a consequence of different amounts of exposed rocks of different ages. The analyses of African and South American records (which had fewer records) were run for 10 million generations, whereas all others were run for 20 million generations. Not all analyses converged but we generally used the results from 10 separate chains, where the effective sample size for all key parameters was high (i.e. all parameters related to the overall process but not necessarily the speciation or extinction time of each species, which are treated as individual parameters, had an effective sample size above 200). The exception for this was Europe, where convergence by these criteria was only seen in four chains and only these four were therefore used in the subsequent analyses.

Finally, we combined the estimated extinction dates from the PyRate analyses with the origination times estimated from the phylogenetic analyses. We first sampled random generations across the different PyRate analyses (with the same number of samples for each). Following this, we combined the results from a random PyRate generation with a random tree from the posterior distribution. By doing this, we estimated the extinction time for every species on one continent independently of their extinction time on other continents, which means that we consider them to represent separate distinct populations rather than meta-populations with ongoing gene flow. We treated Eurasia as a continent for all downstream analyses rather than as the distinct continents of Europe and Asia. In doing this we set the Eurasian extinction date as the latest of the inferred European and Asian extinction dates.

#### 5.2 Data analyses

##### 5.2.1 Dispersal inference

The first analytical step was an estimation of the dispersal dynamics between continents. For this and all further analyses we discarded pinnipeds (which were just included in the datamining elements to maximise the use of our created phylogeny for future evolutionary studies) and thus only analysed terrestrial species. We estimated ancestral geographic range for all nodes with a DEC (dispersal-extinction-cladogenesis) model in BiogeoBEARS (47), using the trees with origination time as described above. We used a DEC rather than the frequently used DEC +j model since the underlying mathematical properties of the DEC +j model have been seriously questioned (48).

We manually specified settings in BiogeoBEARS to match the study system. We did not allow dispersal to South America prior to 10 million years ago (Ma). The oldest carnivore fossils from the continent within our database are two records of the procyonid genus *Amphinasua* dated to 6.8–9 Ma; i.e. the Huayquerian South American Land Mammal Age (SALMA). By doing this we assume that at most one SALMA (the Chasicoan, 9–10 Ma) could lack fossils, even though carnivores actually could be present. We only allowed dispersal between adjoining continents (Africa/Eurasia, Eurasia/North America, and North America/South America). Finally, we allowed the dispersal rate in the Pleistocene (and Holocene) to be potentially higher than the pre-Pleistocene dispersal rate but otherwise kept a single constant dispersal rate. BiogeoBEARS does not generally allow temporal variation in dispersal rates, unless the relative temporal rates are manually specified, but we allowed the dispersal rate in the Pleistocene to be different from the pre-Pleistocene rate by setting the Pleistocene rate as d_Pleistocene_ = d_Pre-Pleistocene_*2*^w^*, with *w* being a free parameter estimated by BiogeoBEARS.

As a second step, we transformed the probabilistic ancestral states at all nodes of the DEC analysis into binary presence/absences by sampling values based on the estimated probabilities. This was done starting with the most terminal nodes. While sampling ancestral nodes, we only sampled among states that were reasonable considering the states of the descendant nodes. Thus, if the estimated ancestral state of two daughter nodes was inferred to be Eurasia and North America, the ancestral area of their direct ancestor was restricted to combinations of one or both of these two areas.

Finally, we estimated dispersal times between continents. Whenever mechanistically plausible (e.g. if the ancestor lived in North America and the daughter species lived in Eurasia and North America) dispersal was inferred to be at the time of speciation. When this was not the case, the necessary dispersal events (and potential required extinction events) were placed equidistant from each other on the relevant branches.

We acknowledge that the procedure of assigning areas to nodes and branches can be seen as a violation from the DEC model the data was estimated under, but we consider these violations biologically justifiable. Our estimation procedure means that we can infer jump dispersal whenever it is possible. These do not exist under a standard DEC model, but due to mathematical problems with the extension that explicitly incorporates jump dispersal (DEC+j model), we preferred to use this workaround. If we instead had used stochastic mapping to infer ancestral areas, we would have drastically overestimated the magnitude of dispersal events – especially ones taking place along long branches. This is because the only way to generate the frequent range changes that can be seen for rapidly diversifying clades without having jump dispersal in the model is by having unrealistically high dispersal rates within lineages. To get an idea of the magnitude of this issue, we estimated ancestral areas through stochastic mapping ten times for each of the 100 trees. We found that the smallest overestimation among all these 1,000 replicates was 47% while the median overestimation was 71%. The overestimation of dispersal events based on stochastic mapping would be particularly problematic for our analyses, due to its concentration on longer branches, which would produce biased results in our analyses of the relationship between dispersal rate and diversification.

##### 5.2.2 Statistical analyses

We conducted a number of separate analyses on 100 separate trees from the posterior distribution. These analyses dealt with the dispersal dynamics between continents, the build-up of diversity and the macro-evolutionary success of intercontinental dispersers relative to other species. In all cases described below, we analysed the patterns in standard regression analyses with the results from each of the 100 trees weighted equally.

Some analyses (related to phylogenetic diversity and diversification rate) are only meaningfully interpretable for ultrametric trees. For simplicity, extinction was therefore dealt with in time intervals and on trees sliced at various ages, rather than in continuous time, counting only the species (internal or external branches) extant at that time point. Hence, when using 0.5-million-year time intervals, two species that went extinct 1.2 and 1.4 Ma would both be included as extant for a tree sliced at 1.0 Ma (but extinct in a tree sliced at 0.5 Ma). All analyses were conducted with time intervals of 0.25, 0.5 or 1.0 million years duration in order to test the effect of this procedure on the results.

In the simplest analyses, we calculated species diversity and phylogenetic diversity (49) (i.e. the sum of the branch lengths in the tree) for all species alive in each time interval globally or on each continent. In two other sets of analyses, we tested whether dispersing species were diversifying faster than others were. The first of these sets of analyses, which we call *pre-dispersal success*, investigated if species that disperse belong to lineages that, at their time of dispersal, diversified faster than the other lineages present at that point in time. The second, which we call *post-dispersal success*, investigated if species that disperse leave more descendant species than species that do not disperse.

We estimated *pre-dispersal success* based on the diversification rate (DR) (45). We sliced the tree at the end of each time interval throughout the Cenozoic and calculated the DR of all lineages alive at that time. For all intercontinental dispersal events occurring in the following time interval, we then identified the lineage representing the disperser (this could be either the same species or one of its ancestors). We then calculated the logarithm of the ratio between the diversification rate of the disperser and the median diversification rate for either all lineages alive in the time interval (*global pre-dispersal success*) or the subset of these that was found on the same continent the disperser originated in (*continental pre-dispersal success*). This calculation is outlined in Fig. 3.

We estimated *post-dispersal success* by comparing the tree at the time before dispersal with a tree sliced a number of million years afterwards. For each lineage alive at the first time interval, we identified how many species they had diversified into a few million years later (this would be 0 if the lineage had gone extinct in the meantime). We then calculated the ratio between the number of species in the dispersing lineage and the mean for either all other species (*global post-dispersal success*) or all species from the continent dispersal is to (*continental post-dispersal success*). In both cases, we square-root transformed the *post-dispersal success* to improve normality. For this measure, we used square-root rather than log transformation and means rather than medians, because zero descendants for both the dispersing and non-dispersing lineages are common. Zero descendants for the dispersing lineages could otherwise require taking the logarithm to zero, while zero descendants for the non-dispersers (if occurring for more than half the species) would otherwise require dividing by zero. Dispersal events were ignored for these analyses if they occurred so recently that the time period a few million years later than that which we compare them to would be in the future. Separate analyses of *post-dispersal success* were conducted on trees sliced after 3, 5, and 7 My (with all dispersal events occurring within the last 3, 5, or 7 My ignored).

##### 5.2.3 Simulations

All analyses described in 5.2.2 implicitly assume complete sampling. Although we have included all known species, an unknown number of extinct species may be missing from the fossil record, which can influence our results. This is especially so since the fraction of missing species is expected to vary in time and space. In order to understand the influence of missing taxa on our results, we, therefore, simulated a number of random phylogenies. We then simulated incomplete sampling on those phylogenies, then repeated all analyses described in 5.2.2 on both the full and the sampled phylogenies.

We simulated trees based on a stage-dependent speciation and extinction model. More specifically, we simulated trees based on a seven-class ClaSSE (Cladogenetic State change Speciation and Extinction) model (61) modified into a four-area version of the normally two-area GeoSSE model (62) using Diversitree (63). In this version, each species was given seven potential character states 1: S, 2; SN, 3: N, 4: NE, 5: E, 6: EA, 7: A (where S means South America, N means North America, E means Eurasia and A means Africa). The model included five parameters: sympatric speciation rate (λ1) present in all classes; jump dispersal speciation (λ2) for single area classes; allopatric speciation (λ3) for two area classes; local extinction rate (ε); and dispersal rate (δ) only between adjoining regions. The model is outlined in Fig. S2.

We generated plausible trees of the same size as the empirical ones using a rejection sampler to obtain a distribution of trees resembling the empirical ones in shape and geographic ranges. We obtained the ClaSSE parameter values by randomly drawing them from the following uniform distributions: λ1 ∼ U(0, 0.2), λ2 ∼ U(0,0.05), λ3 ∼ 3 ∼ U(0,3), δ ∼ U(0, 0.1), and ε ∼ U(0, 0.4). We generated phylogenetic trees and geographic ranges at each random draw. The rejection sampler included three summary statistics: 1) the fraction of all taxa that are extant; 2) the total number of dispersals; and 3) the number of extant species occurring on more than one continent. Phylogenetic trees and geographic ranges were only accepted if all summary statistics met the condition: 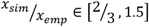 where x_sim_ is the statistic obtained from the simulation (e.g. the total number of dispersals) and x_emp_ is the corresponding value obtained from the empirical data. We repeated the simulation until 100 trees were accepted. The branch lengths of these trees were then multiplied by the appropriate factors to have root ages corresponding to those of the 100 empirical trees.

We simulated geographic and spatial variation in preservation rate for each of these random trees based on estimated sample intensity at each epoch as inferred from the PyRate analysis (see 4.1.4). We assumed complete sampling of extant species. Sample intensity for the PyRate analysis was estimated separately for Asia and Europe, due to the large difference in paleontological research conducted in the two continents. For all biogeographical analyses, however, we used a combined Eurasia since the borders between the two are poorly defined and a large fraction of species have ranges spanning both continents. For the simulations (which are intended to mimic the biogeographical analyses), we, therefore, used a combined value for Eurasian sampling. This was estimated as the mean of the European and Asian sampling weighted by the contemporary diversity of carnivores in the two continents. No pre-Miocene South American carnivores exist in the empirical trees and therefore we cannot use empirical values for this continent for the Paleocene, Eocene, and Oligocene. We instead used the mean estimate for Africa and Eurasia (corresponding to substantially lower estimated sampling effort than North America). Estimation of sample intensity in PyRate can be imprecise for very shallow time intervals such as the Holocene (Daniele Silvestro, pers. comm.). We, therefore, used the Pleistocene value for both the Pleistocene and Holocene.

We carried out one round of random sampling based on the preservation rate defined above for each continent, separately for each branch. We first assessed sampling on all external branches on all continents and accepted presence whenever sampling was simulated to have taken place. After this, we assessed internal branches ranked in order of increasing number of descendants. Whenever an internal branch occurred on, and was sampled in, a continent where none of its occurring descendants were sampled, we considered a random descendant species on the relevant continent as sampled instead. The logic of this treatment can be understood by looking at a small clade of two species, with a long internal branch and an extinction of both species nearly immediately after speciation. In such cases, the probability of sampling both species would be limited but it is more likely that we would sample the lineage before speciation. If we only looked at sampling in external branches, we would thus drastically underestimate the diversity resulting from incomplete sampling.

In order to test the importance of incomplete sampling, we repeated all analyses from 5.2.2 on both the full random trees and the random trees with simulated sampling. After this, we assessed if the effects we observed in the empirical trees matched the differences between the simulated trees with incomplete and complete sampling, in which case extreme care would be needed in the interpretation of the results.

---

**Table S1:** Continental pre-dispersal success. The test statistic is the logarithm of the ratio between the diversification rate at the time of dispersal of dispersers and the median of the diversification rates of all species present in the source continent just before the dispersal event (see Fig. 1). The p-values for the global rate and for the temporal age effect are the probability of being different from 0. For models with different patterns depending on the source and or target continent, the p-value is based on the probability of being different from the estimated global rate.

|  | 1.0-million-year<br>intervals | 0.5-million-year<br>intervals | 0.25-million-year<br>intervals |
| --- | --- | --- | --- |
| Simplest model |  |  |  |
| Global | 0.0900 (0.0199)<sup>***</sup> | 0.0835 (0.0194)<sup>***</sup> | 0.0785 (0.0190)<sup>***</sup> |
| AIC | 126.202 | 120.325 | 122.352 |
| Temporal variation |  |  |  |
| Global | 0.1229 (0.0260)<sup>***</sup> | 0.1122 (0.0259)<sup>***</sup> | 0.1068 (0.0255)<sup>***</sup> |
| Age | -0.0021 (0.0012)<sup>‘</sup> | -0.0018 (0.0012)<sup>‘</sup> | -0.0018 (0.0012)<sup>‘</sup> |
| AIC | 123.159 | 118.431 | 120.740 |
| Variation between source continent |  |  |  |
| From Africa | 0.0519 (0.0599) | 0.0487 (0.0577) | 0.0422 (0.0570) |
| From Eurasia | 0.1287 (0.0260) | 0.1226 (0.0260) | 0.1185 (0.0253) |
| From North America | 0.0386 (0.0331) | 0.0303 (0.0330) | 0.0240 (0.0329) |
| From South America | 0.1052 (0.1528) | 0.1002 (0.1439) | 0.0980 (0.1407) |
| AIC | 121.666 | 115.841 | 114.379 |
| Variation between target continent |  |  |  |
| To Africa | 0.1191 (0.0374) | 0.1160 (0.0373) | 0.1137 (0.0366) |
| To Eurasia | 0.0488 (0.0340) | 0.0453 (0.0343) | 0.0379 (0.0337) |
| To North America | 0.1351 (0.0356) | 0.1260 (0.0360) | 0.1214 (0.0346) |
| To South America | 0.0192 (0.0513) | 0.0017 (0.0503) | -0.0009 (0.0495) |
| AIC | 121.745 | 114.352 | 114.924 |
| Variation between source and target continent |  |  |  |
| Africa to Eurasia | 0.0519 (0.0600) | 0.0487 (0.0577) | 0.0422 (0.0570) |
| Eurasia to Africa | 0.1191 (0.0374) | 0.1161 (0.0373) | 0.1137 (0.0366) |
| Eurasia to North America | 0.1372 (0.0368) | 0.1281 (0.0370) | 0.1231 (0.0357) |
| North America to Eurasia | 0.0479 (0.0420) | 0.0439 (0.0428) | 0.0362 (0.0429) |
| North to South America | 0.0192 (0.0513) | 0.0018 (0.0503) | -0.0009 (0.0496) |
| South to North America | 0.1051 (0.1529) | 0.1002 (0.1439) | 0.098 (0.1407) |
| AIC | 124.174 | 116.893 | 117.311 |

<sup>‘</sup> 0.10>P>0.05 \* 0.05>P>0.01 \*\* 0.01>P>0.001 \*\*\* 0.001>P

**Table S2:**
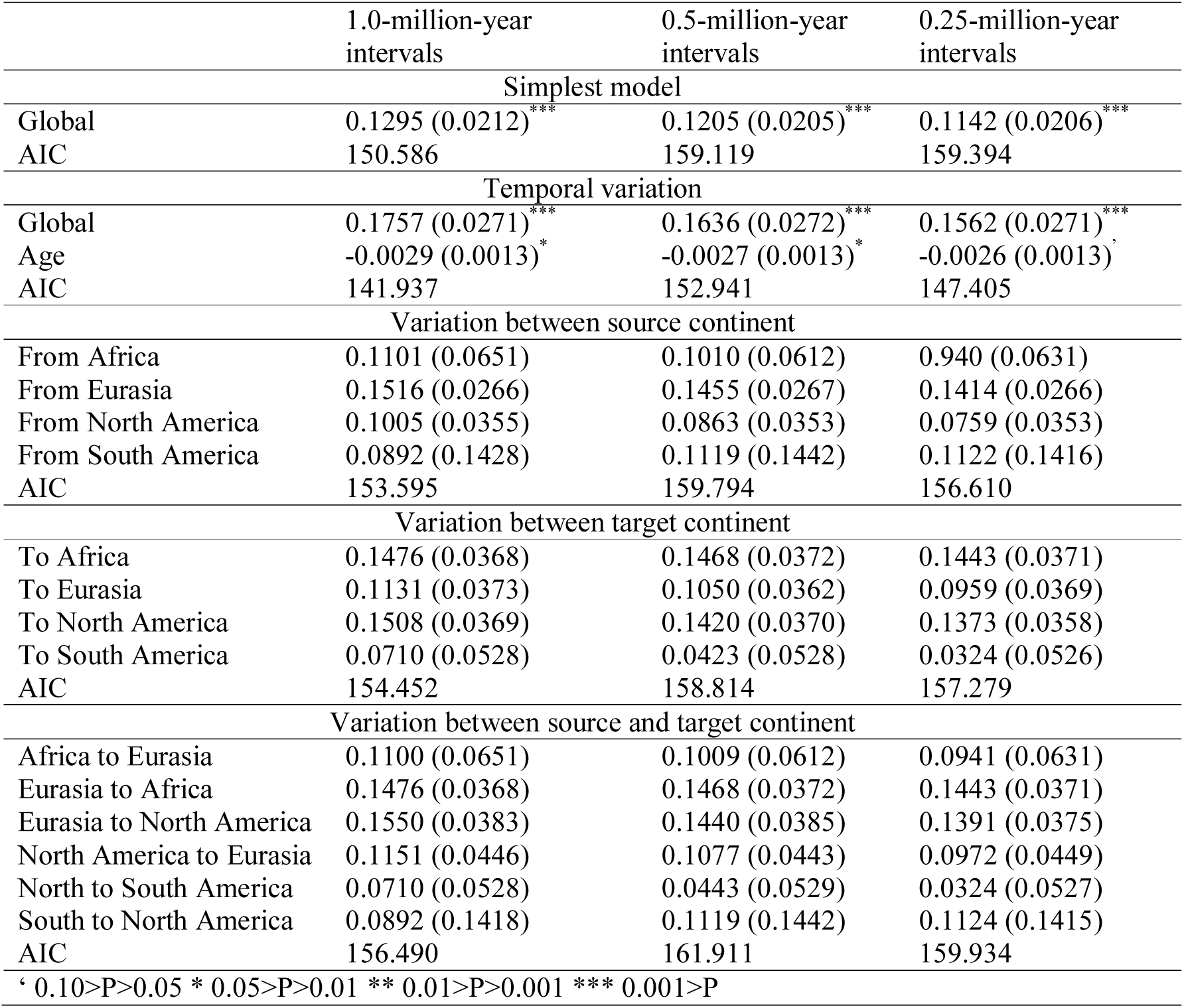
Global pre-dispersal success. The test statistic is the logarithm of the ratio between the diversification rate at the time of dispersal of dispersers and the median of the diversification rates of all species alive globally just before the dispersal event (see Fig. 1). The p-values for global rate and for the temporal age effect are the probability of being different from 0. For models with different patterns depending on the source and/ or target continent, the p-value is based on the probability of being different from the estimated global rate.

**Table S3:**
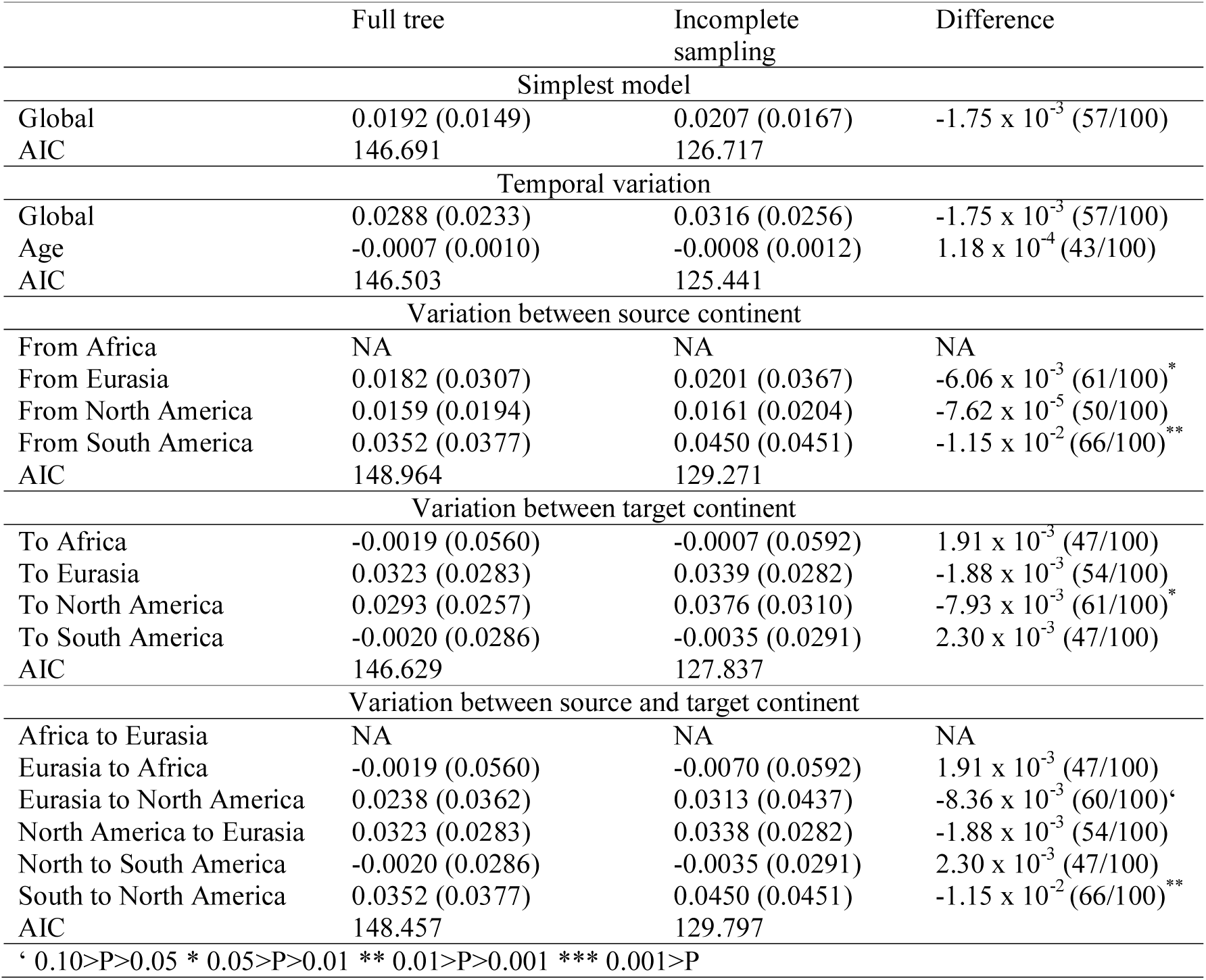
Continental pre-dispersal success in simulations. Values are given for both the full simulated tree and the simulated incomplete sampling. The difference is the median difference between estimates for each tree. In parentheses, we list the number of times where this difference is negative (i.e. how many times the value is larger for incomplete sampling than the full tree). The test statistic is the logarithm of the ratio between the diversification rate at the time of dispersal of dispersers and the median of the diversification rates of all species in the source continent from just before the dispersal event (see Fig. 1). For models with different patterns based on the source and/ or target continent, the p-value is based on the probability of being different from the estimated global rate. The p-value for the difference is based on a two-tailed binomial distribution and tests if incomplete sampling is equally likely to lead to larger and smaller values than complete sampling.

**Table S4:**
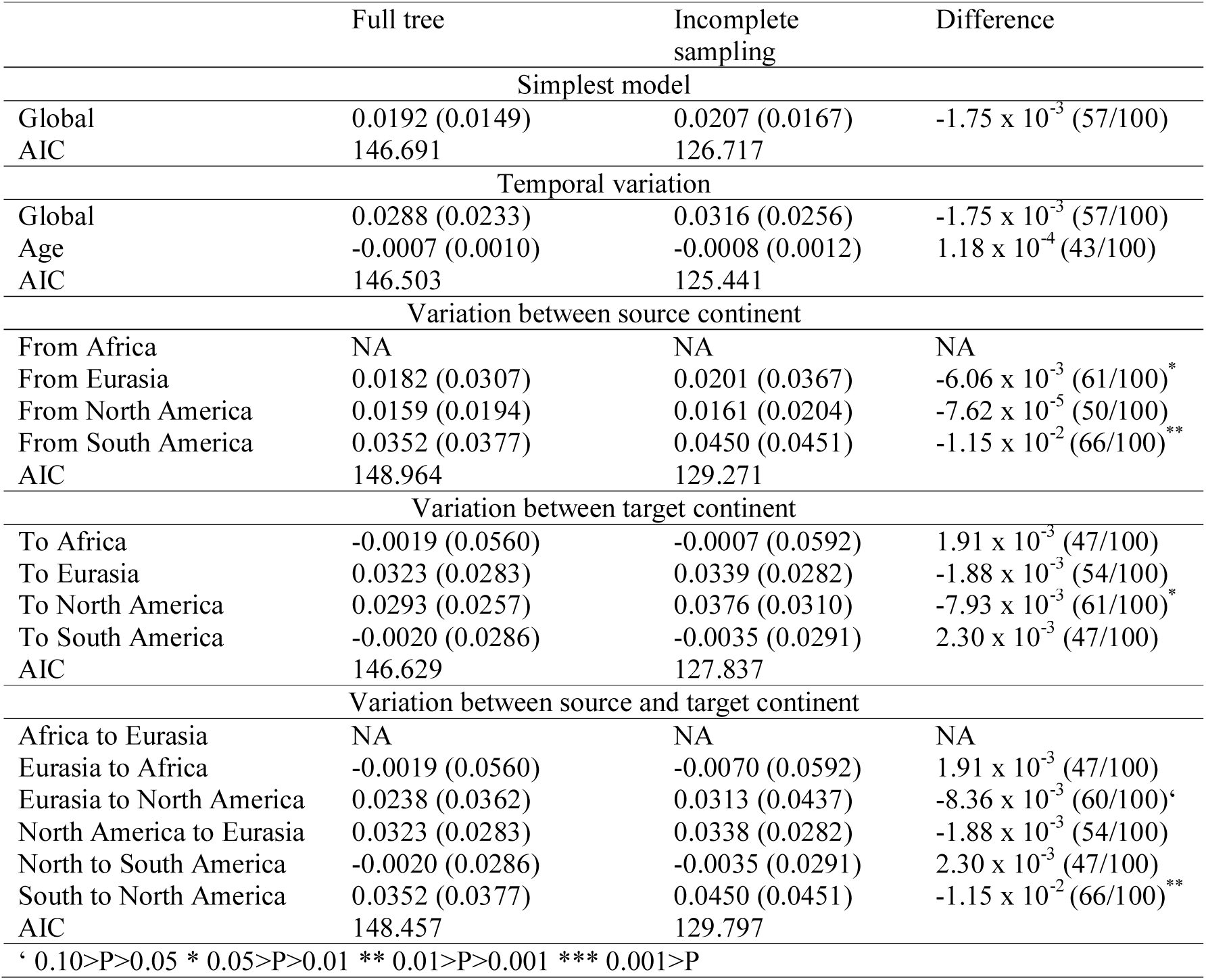
*Global pre-dispersal success* in simulations. Values are given for both the full simulated tree and the simulated incomplete sampling. The difference is the median difference between estimates for each tree. In parentheses, we list the number of times where this difference is negative (i.e. how many times the value is larger for incomplete sampling than the full tree). The test statistic is the logarithm of the ratio between the diversification rate at the time of dispersal of dispersers and the median diversification rates of all species alive globally just before the dispersal event (see Fig. 1). For models with different patterns based on the source and/ or target continent, the p-value is based on the probability of being different from the estimated global rate. The p-value for the difference is based on a two-tailed binomial distribution and tests if incomplete sampling is equally likely to lead to larger and smaller values than complete sampling.

**Table S5:**
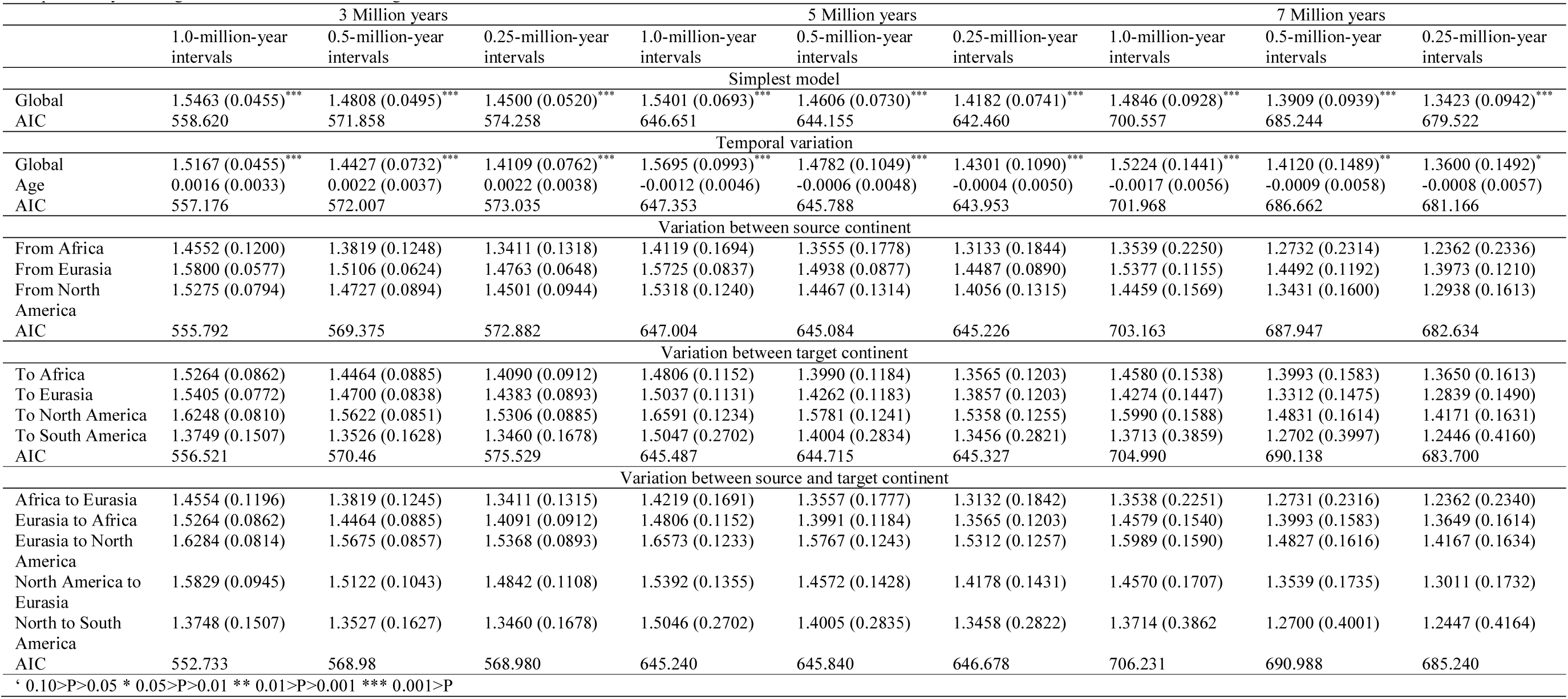
Global post-dispersal success. The test statistic is the square root of the mean number of species alive after a given number of million years (3, 5 or 7) after dispersal, divided by the mean for all species globally. Significance for the estimate is the probability of being different from 1 (the random expectation), for age it is the probability of being different from 0, and for models with different patterns based on the source and/ or target continent, it is based on the probability of being different from the estimated global rate.

**Table S6:**
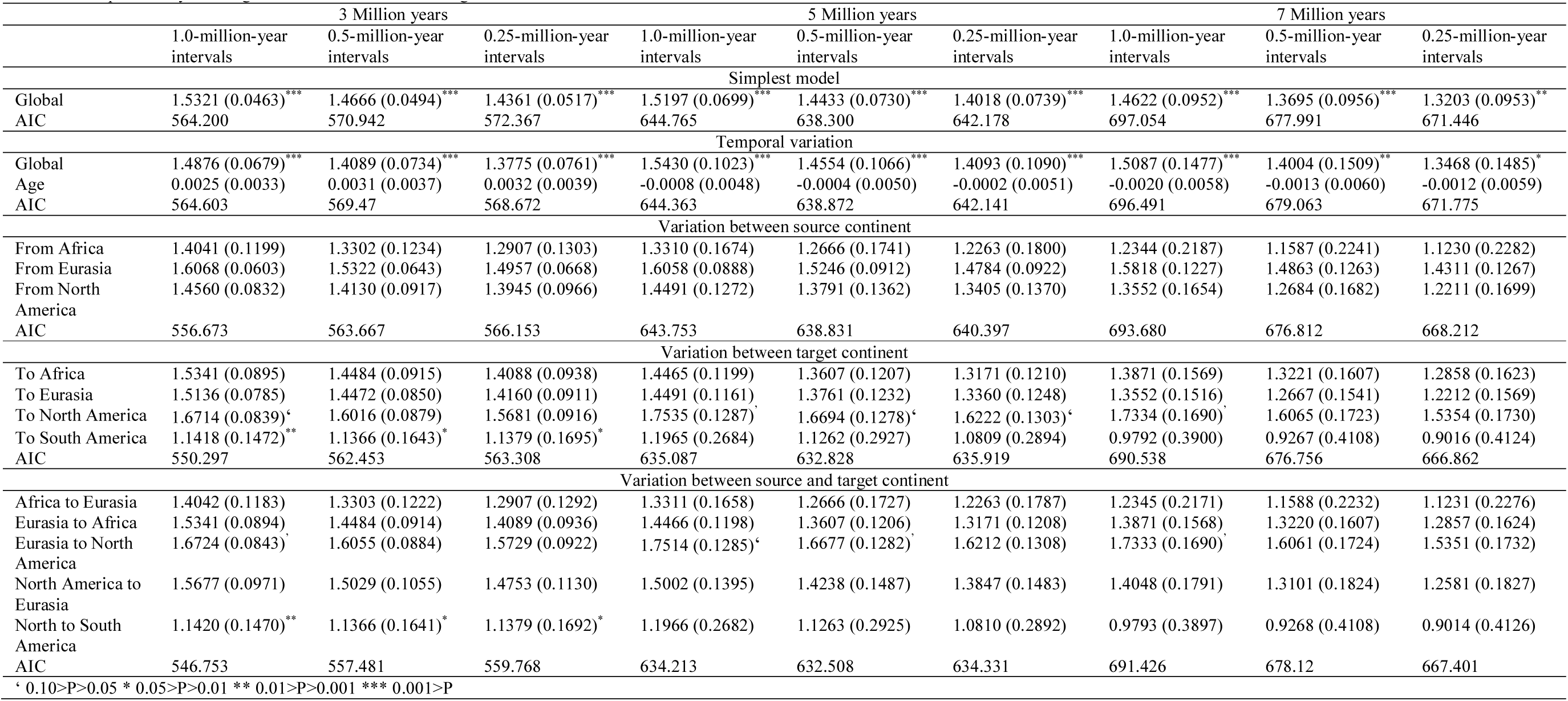
Continental post-dispersal success. The test statistic is the square root of the mean number of species alive after a given number of million years (3, 5 or 7) after dispersal, divided by the mean for species in the target continent. Significance for the estimate is the probability of being different from 1 (the random expectation), for age it is the probability of being different from 0 and for models with different patterns based on target and/ or source continent it is based on the probability of being different from the estimated global rate.

**Table S7:**
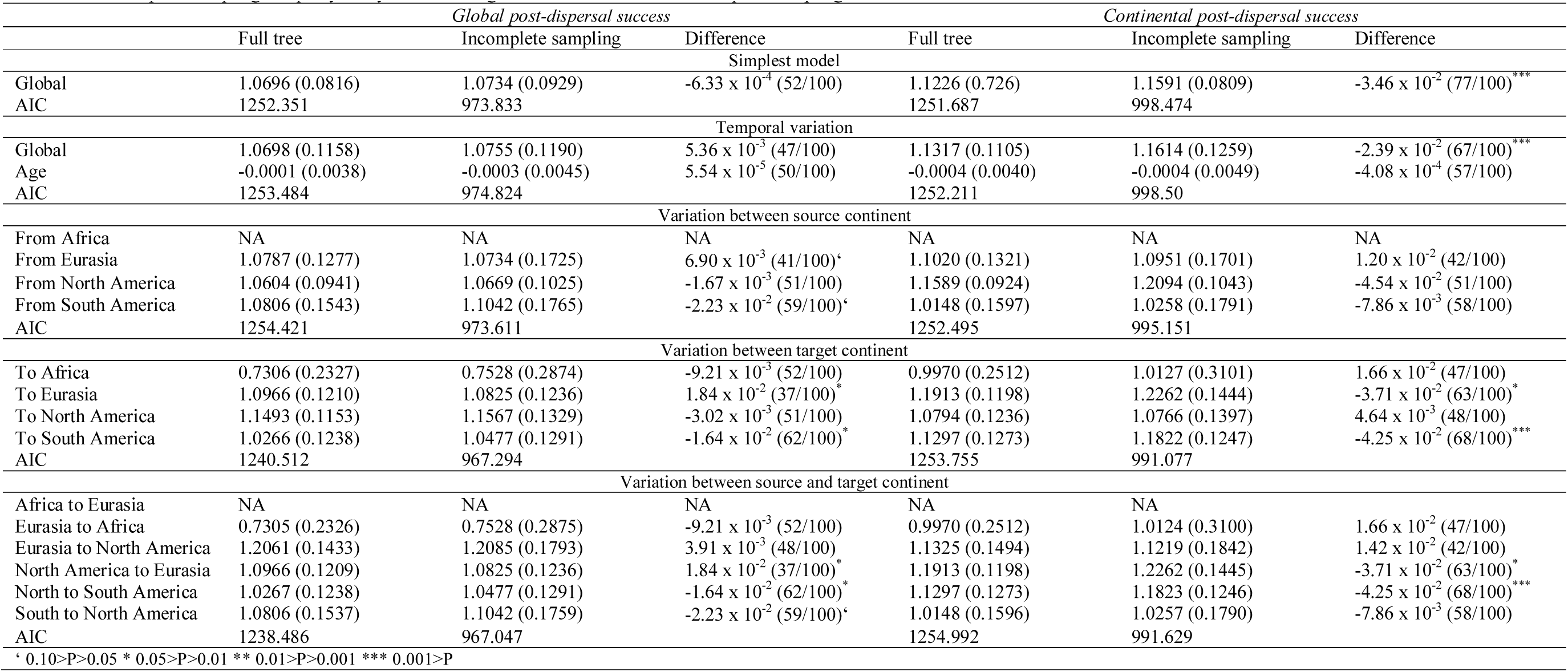
Post-dispersal success in simulations. The test statistic is the square root of the mean number of species alive after 5 million years after dispersal divided by the mean value for all other species globally. Significance for the estimate is the probability of being different from 1 (the random expectation), for age it is the probability of being different from 0 and for models with different patterns based on the source and/ or target continent it is based on the probability of being different from the estimated global rate. Values are given for both the full simulated tree and the simulated incomplete sampling. The difference is the median difference between estimates for each tree. In parentheses, we list the number of times where this difference is negative (i.e. how many times the value is larger for incomplete sampling than the full tree). The p-value for the difference is based on a two-tailed binomial distribution and tests if incomplete sampling is equally likely to lead to larger and smaller values than complete sampling.

**Table S8:**
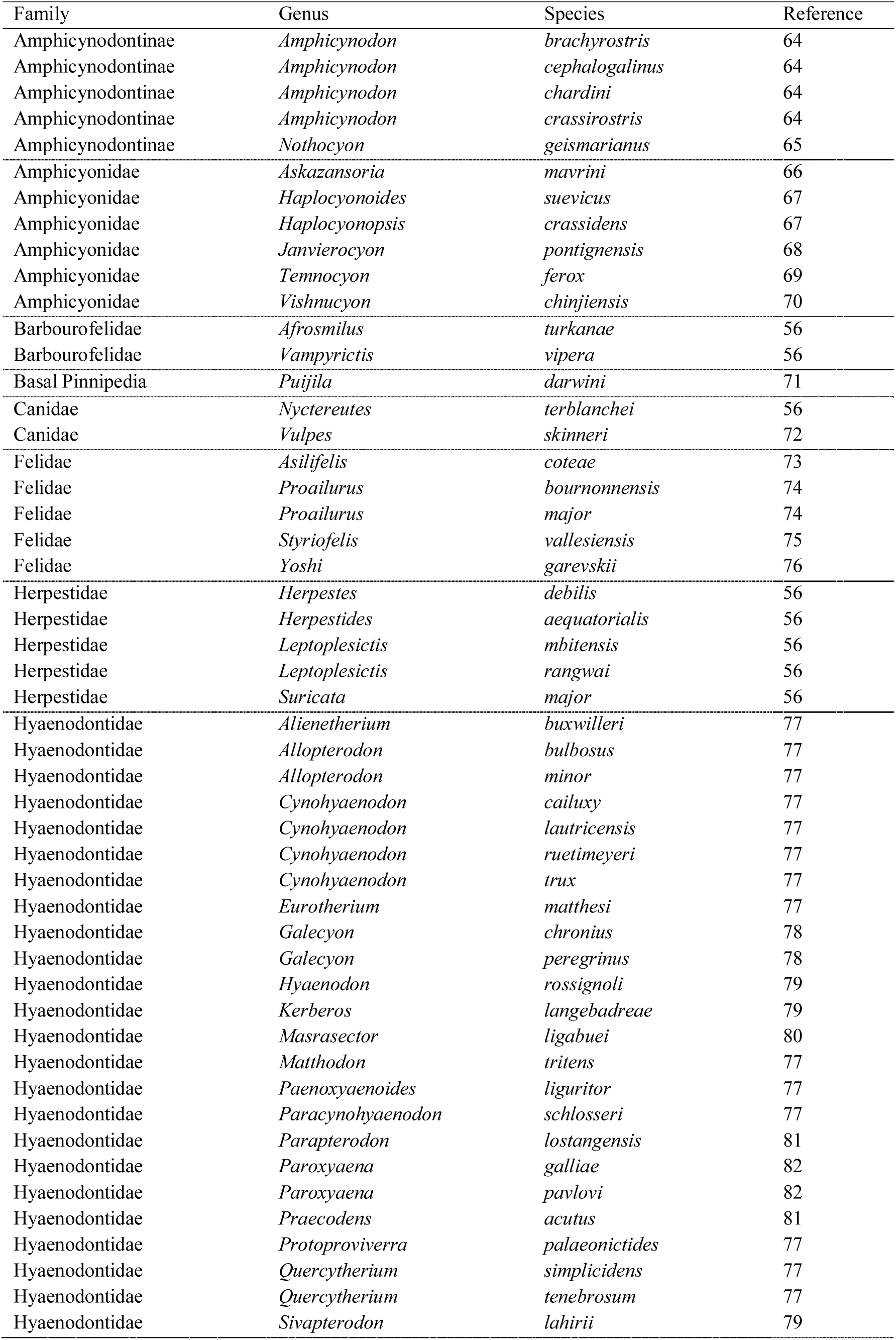
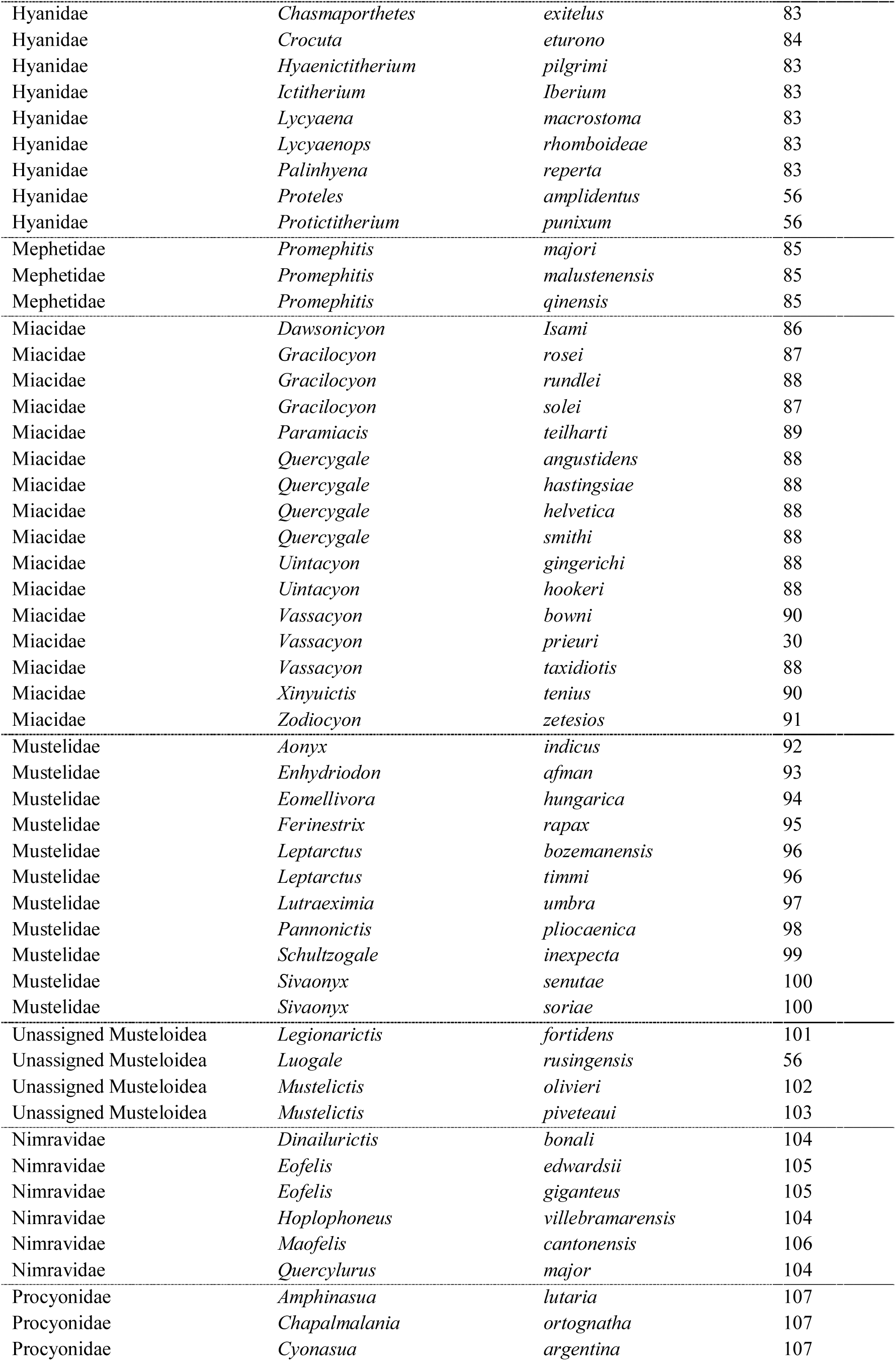
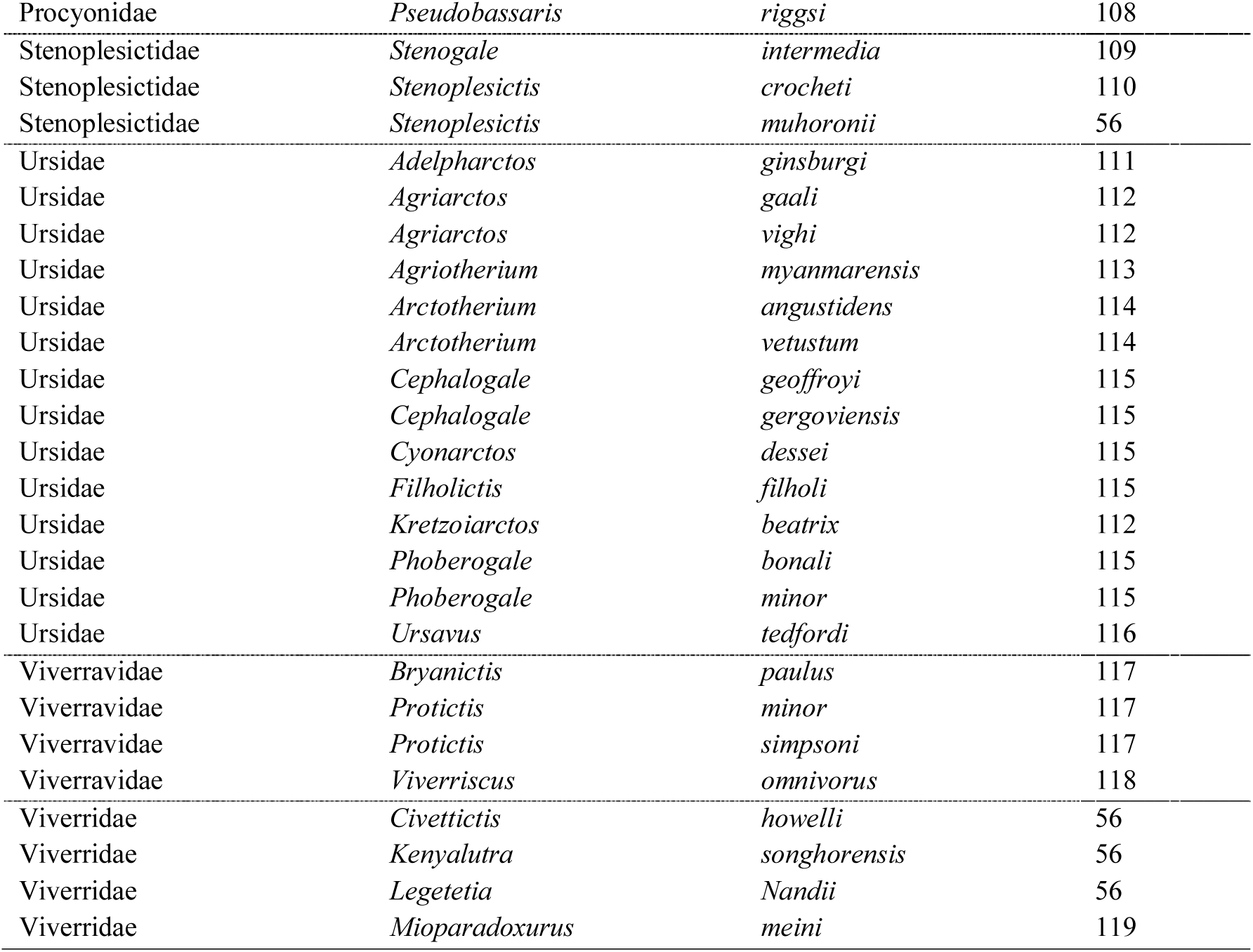
Additional species not in NOW or PBDB.

**Table S9:**
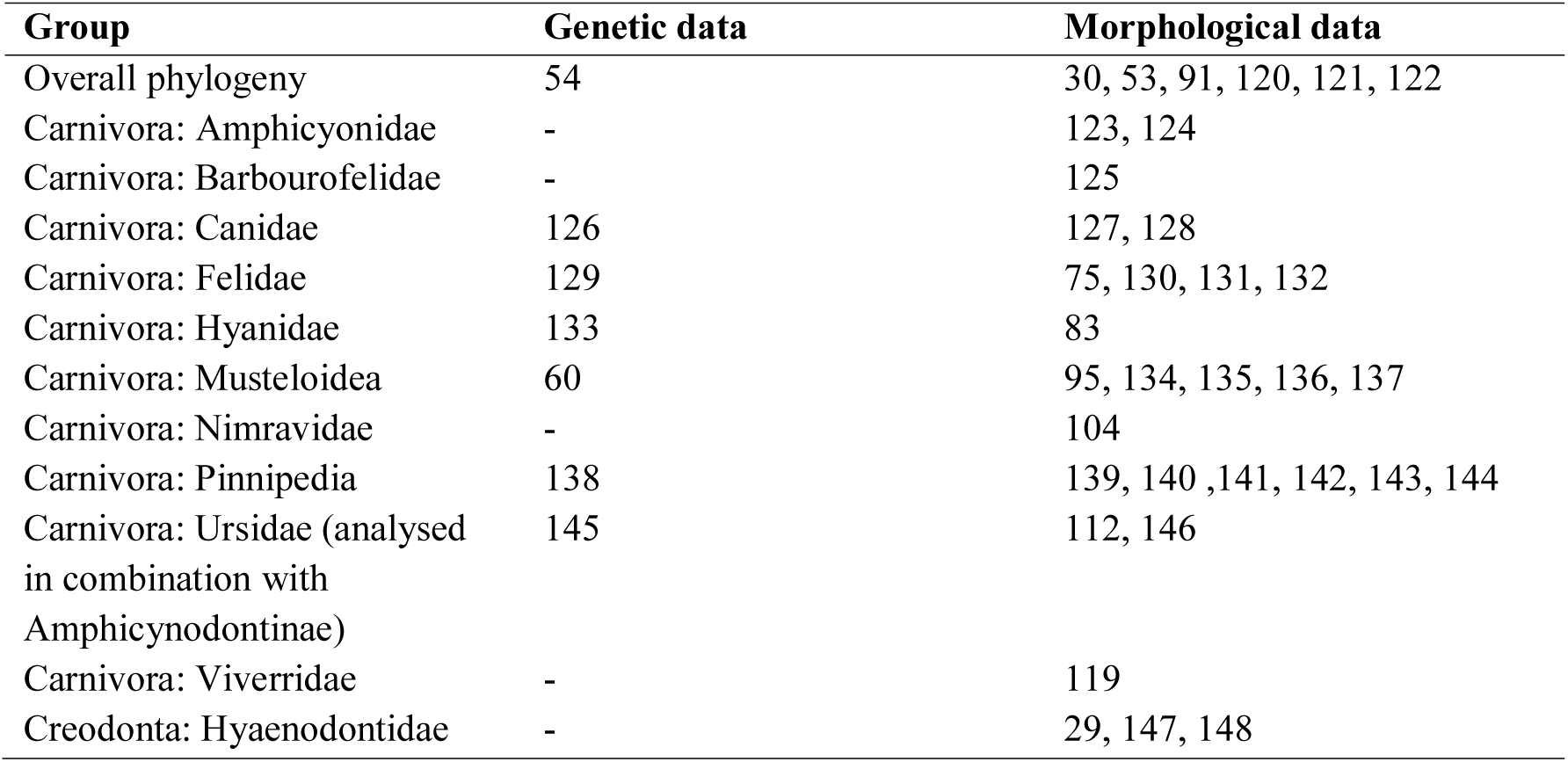
Sources for morphological and genetic data within groups.

**Table S10:**
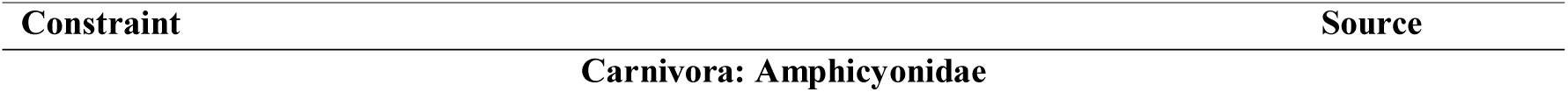
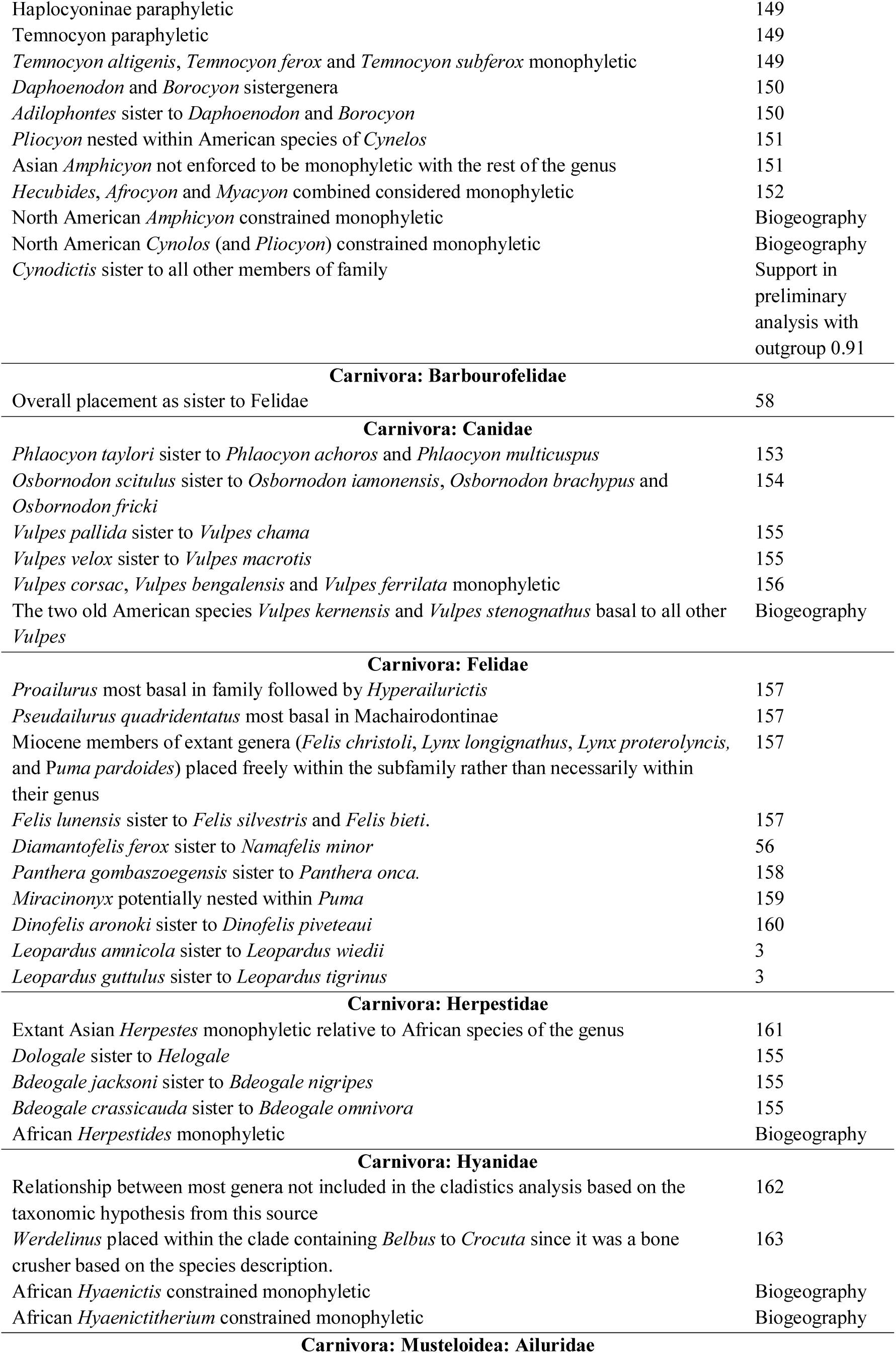
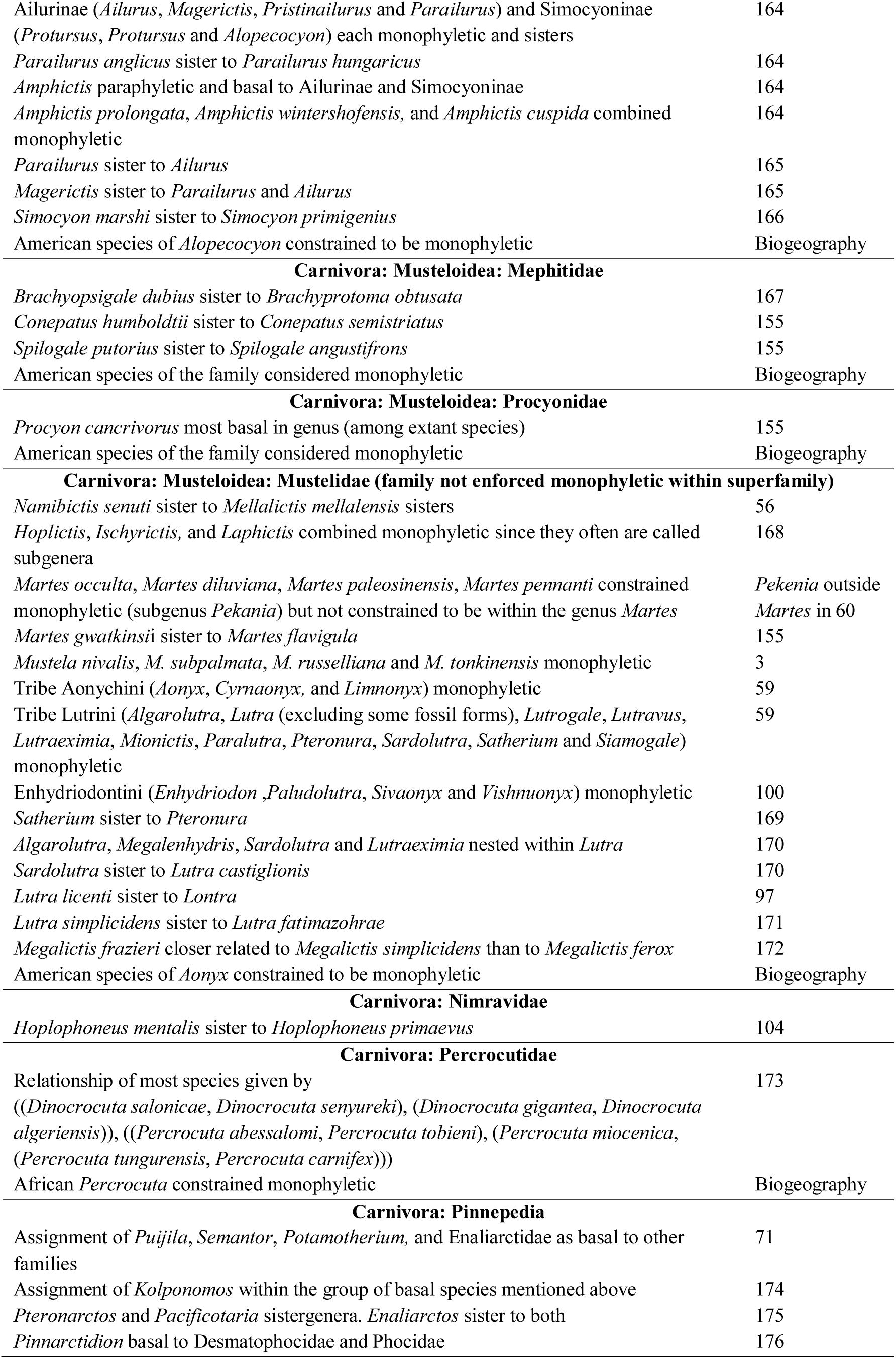
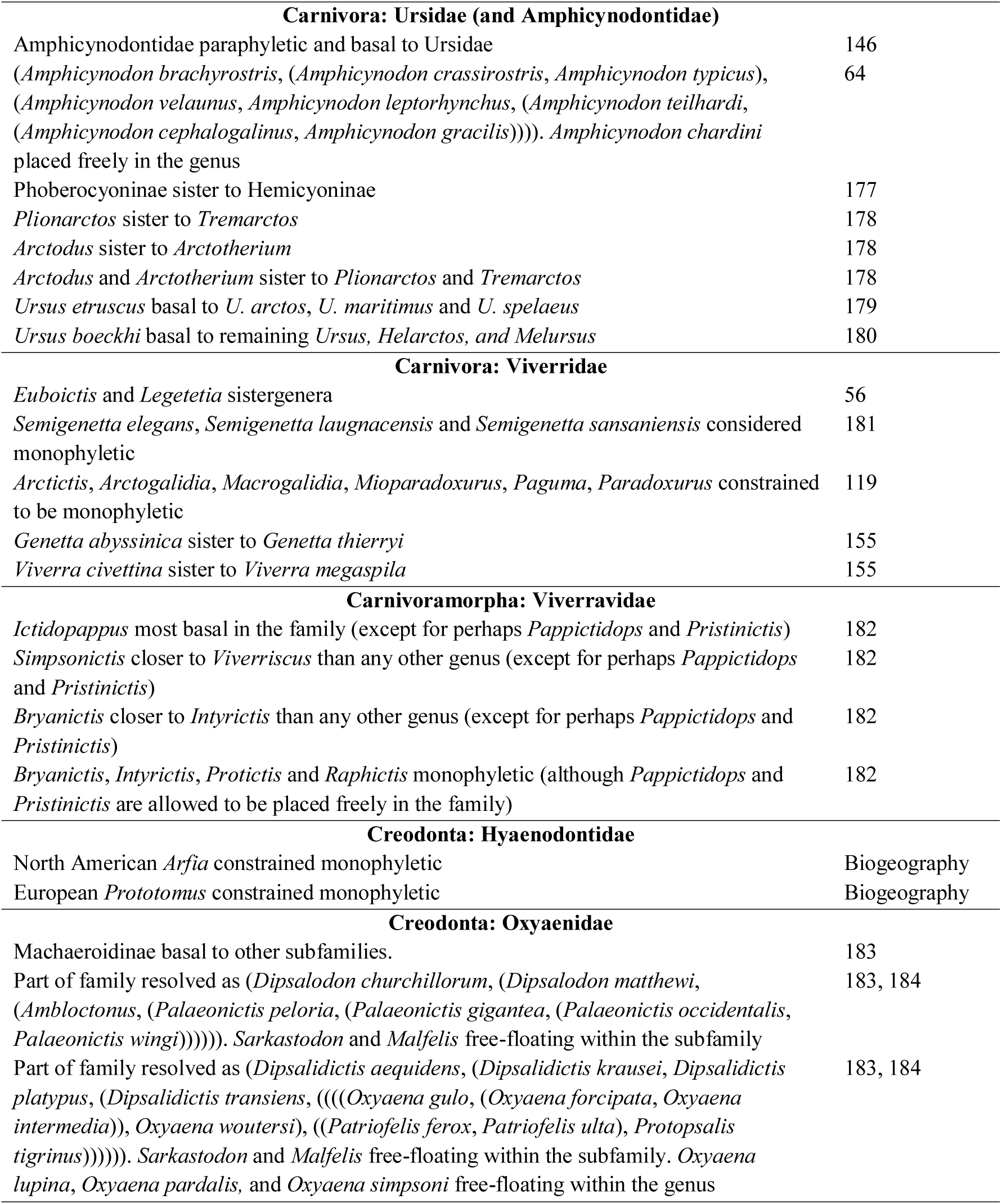
Topological constraints.

**Figure S1:**
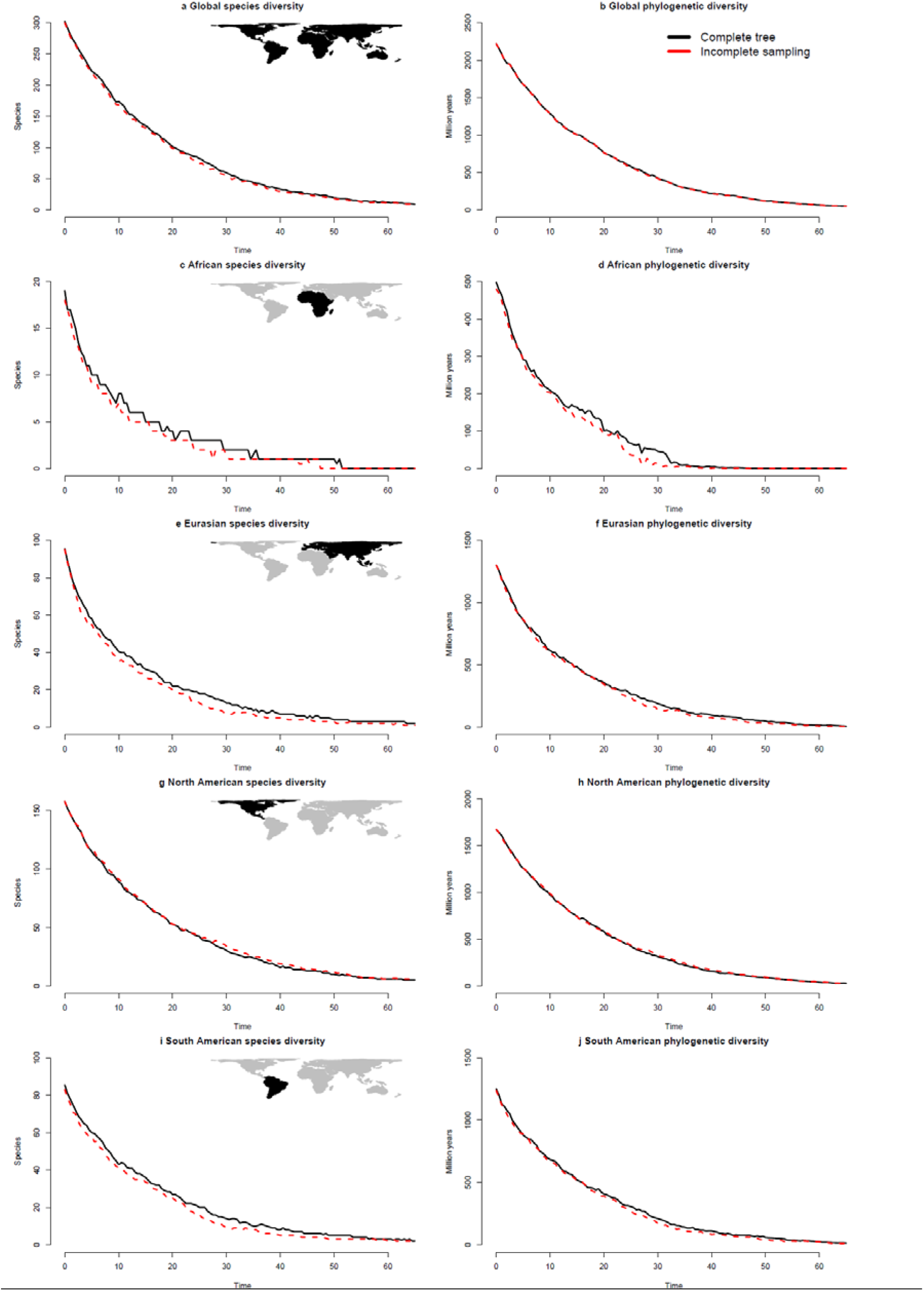
Effect of incomplete sampling on diversity estimates. The difference in median species and phylogenetic diversity either globally or for individual continents. Due to the limited effect of limited sampling, the two lines are frequently on top of each other and the line for incomplete sampling is therefore stippled to make both lines visible.

**Figure S2:**
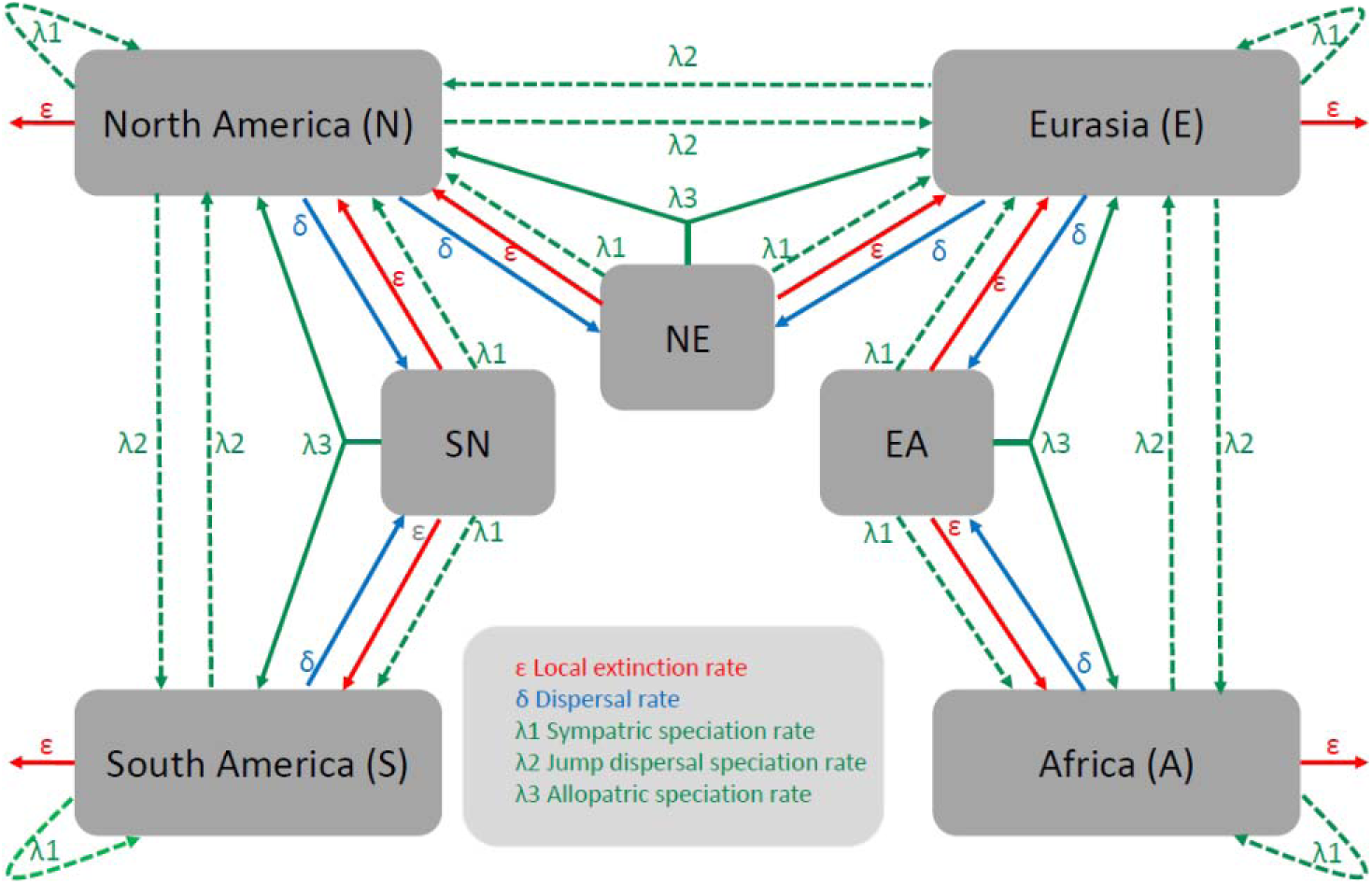
Simulation model. Schematic drawing of the simulation model, which is a seven classes ClaSSE model transformed into a four-area version of a GeoSSE model. All arrows drawn in full (dispersal rate, extinction rate and allopatric speciation rate) represent cases where the species in question changes class in the ClaSSE model (or completely disappears for extinction from single area classes), whereas stippled lines (sympatric speciation rate and jump dispersal speciation rate) represent cases where the species buds off from another species while the ancestor stays in the region it was before.

